# Thermodynamic limitations on brain oxygen metabolism: physiological implications

**DOI:** 10.1101/2023.01.06.522942

**Authors:** Richard B. Buxton

**Affiliations:** Center for Functional Magnetic Resonance Imaging, Department of Radiology, University of California San Diego

**Keywords:** cerebral blood flow (CBF), cerebral metabolic rate of oxygen (CMRO_2_), brain energy metabolism, thermodynamic limits on brain function

## Abstract

A recent hypothesis is that maintaining the brain tissue ratio of O_2_ to CO_2_ is critical for preserving the entropy increase available from oxidative metabolism of glucose, with a fall of that available entropy leading to a reduction of the phosphorylation potential and impairment of brain energy metabolism. The hypothesis suggests that physiological responses under different conditions can be understood as preserving tissue O_2_/CO_2_. To test this idea, a mathematical model of O_2_ and CO_2_ transport was used to calculate how well different physiological responses maintain tissue O_2_/CO_2_, showing good agreement with reported experimental measurements for increased neural activity, hypercapnia and hypoxia. The results highlight the importance of thinking about brain blood flow as a way to modulate tissue O_2_/CO_2_, rather than simply in terms of O_2_ delivery to the capillary bed. The hypoxia modeling focused on humans at high altitude, including acclimatized lowlanders and adapted populations, with a primary finding that decreasing CO_2_ by increasing ventilation rate is much more effective for preserving tissue O_2_/CO_2_ than increasing blood hemoglobin content. The modeling provides a new framework and perspective for understanding how blood flow and other physiological factors support energy metabolism in the brain under a wide range of conditions.

**Key points summary:**

- Recent thermodynamic modeling suggests that preserving the O_2_/CO_2_ ratio in brain tissue is critical for preserving the entropy change available from the oxidative metabolism of glucose and the phosphorylation potential underlying energy metabolism.
- The hypothesis tested is that normal physiological responses (notably blood flow changes) often act to preserve this ratio under changing conditions.
- Using a detailed model to calculate tissue O_2_/CO_2_ we found good agreement with the predictions of the hypothesis and reported experimental results during hypoxia, hypercapnia and increased oxygen metabolic rate in response to increased neural activity.
- For the hypoxia modeling we considered high altitude acclimatization and adaptation in humans, showing the critical role of reducing CO_2_ in preserving tissue O_2_/CO_2_.
- The tissue O_2_/CO_2_ hypothesis provides a useful perspective for understanding the function of observed physiological responses under different conditions in terms of preserving brain energy metabolism, although the mechanisms underlying these functions are not well understood.

## 1. Introduction

In an earlier paper we proposed a hypothesis for limitations on oxygen metabolism in the brain based on a thermodynamic effect (1). The essential hypothesis is that the ratio of O_2_ concentration to CO_2_ concentration in tissue (abbreviated as tissue O_2_/CO_2_) is a critical parameter, and that normal physiology often acts to preserve this ratio under changing conditions. For example, the increase in the cerebral metabolic rate of oxygen (CMRO_2_) when neural activity increases would decrease tissue O_2_ if nothing else changed. Cerebral blood flow (CBF) can be viewed as a mechanism to increase tissue O_2_, and based on a simple model of O_2_ transport we calculated how large the CBF change needs to be for a given change in CMRO_2_ (1). The result of the modeling was that to preserve the baseline value of tissue O_2_/CO_2_ requires that the fractional CBF increase needs to be 2-3 times higher than the fractional increase of CMRO_2_. An apparent mismatch of CBF and CMRO_2_ responses to increased neural activity in this range has been observed in many studies of neural activation, suggesting that the CBF response to increased neural activity is approximately maintaining tissue O_2_/CO_2_. This imbalance of the change in CBF and CMRO_2_ is the origin of the blood oxygenation level dependent (BOLD) effect, the basis for functional MRI to map changes in neural activity (2).

In the earlier work (1) the initial demonstrations of the implications of the hypothesis were based on a simplified model of O_2_ and CO_2_ transport in blood to estimate the tissue O_2_/CO_2_ ratio. The current work has two goals: 1) To expand the modeling with a more complete treatment of factors affecting O_2_ and CO_2_ transport to improve the estimates of tissue O_2_/CO_2_; and 2) To use the modeling to analyze the implications of the hypothesis for the dynamics of CBF and CMRO_2_ in hypoxia. For the latter, our central question was: in coping with hypoxia, which physiological strategies are most effective for maintaining tissue O_2_/CO_2_ over the normal range of variation of CMRO_2_ associated with brain function? Here we focused on two strategies: increasing the ventilation rate to lower CO_2_, and increasing blood hemoglobin concentration to maintain O_2_ delivery to the tissue capillary bed.

As examples seen in nature, human populations at high altitude potentially illustrate different strategies for coping with hypoxia: 1) lowlanders acclimatized to high altitude after several weeks of exposure; 2) Andean highlanders, who are estimated to have adapted to living at high altitude over the last 11,000 years; and 3) Tibetan highlanders, who are estimated to have adapted to living at high altitude over the last 25,000 years (3). Our goal was to use the modeling to ask: how well are each of these groups preserving brain tissue O_2_/CO_2_, both for normal levels of CMRO_2_ and for increased CMRO_2_ associated with neural activity?

## 2. Theory: the O_2_/CO_2_ hypothesis

The O_2_/CO_2_ hypothesis is that preserving the tissue ratio of O_2_/CO_2_ is important for preserving the positive entropy change available from oxidative metabolism, and if this falls too much the entropy available from ATP to drive cellular work also degrades. The key ideas are summarized in the following sections and developed in more detail in (1, 4).

### 2.1 The role of entropy changes

Nearly all cellular work ultimately depends on harnessing a large positive entropy change from the conversion of ATP to ADP to drive thermodynamically uphill processes. Importantly, though, it is not the absolute amount of ATP present that is critical, but rather the ratio of ATP to ADP. The entropy change (Δ S_ATP_) associated with the conversion of one ATP molecule to one ADP molecule plus P_i_ can be written as (1):

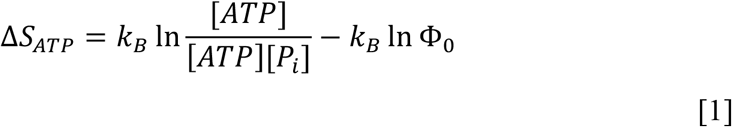

In this equation, brackets indicate concentration and *k*_*B*_ is Boltzmann’s constant. The ratio in the first term is called the ‘phosphorylation potential’, and Φ_0_ is the value of that ratio at equilibrium. If the phosphorylation potential in the cell is equal to Φ_0_, there is no entropy change when ATP is converted to ADP. The utility of ATP as a source of a positive entropy change depends on maintaining the ATP/ADP concentration ratio much higher than the equilibrium value. We defined ΔS_ATP_ as the entropy change of 1 ATP converted to 1 ADP, and the entropy change for the reverse process, converting 1 ADP back to 1 ATP, is just -ΔS_ATP_. (The same equation applies to the reverse transformation, but now with the ratios in the logarithms inverted, which simply multiplies the original expression by negative one.)

After ATP is consumed in cellular work in the brain, the ATP is restored by coupling the conversion of ADP back to ATP with the oxidative metabolism of glucose. The glucose is first converted to 2 pyruvate molecules in the cytosol, coupled to the synthesis of 2 ATP from ADP. The bulk of the ATP production, though, happens in the mitochondria, where 1 pyruvate combines with 3 O_2_ to form 3 CO_2_ coupled to pumping protons against their gradient across the mitochondrial inner membrane (an uphill thermodynamic process). The movement of protons back down their gradient (thermodynamically downhill) then drives the conversion of another 17 ADP to 17 ATP. The net result (including the cytosol generated ATP) is 36 ATP generated for oxidative metabolism of 1 glucose and 6 O_2_.

Focusing now just on the activity in the mitochondria, we can define ΔS_OX_ as the entropy change for the conversion of 1 pyruvate plus 3 O_2_ to 3 CO_2_, and write it as in Eq [1] as:

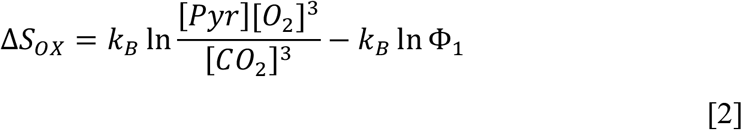

Here Φ_1_ is the equilibrium value of the ratio in the first term.

Eqs [1-2] are expressed in terms of the entropy change, although it is more common to express the same result in terms of a free energy change. The basic idea is that an entropy change, ΔS, can be expressed as being equivalent to the entropy change that happens when an amount of energy ΔE is dissipated as thermal molecular motions in a thermal bath with temperature T: ΔS = ΔE/T. The free energy change is defined as ΔG = -ΔE (i.e., when ΔS is positive ΔG is negative). Instead of working with the entropy change of one transformation (e.g., metabolism of one pyruvate molecule, which is useful for our development below), it is more common to define ΔG for transformation of a mole of molecules, with the gas constant *R* equal to *k*_*B*_ times the number of molecules in one mole (as in (5)). Here we retain the more general ΔS description because it meshes with the usual formulation of the Fluctuation Theorem (6), used in the following section.

### 2.2 Dependence of reaction rates on the entropy change

For physiological entropy transformations, the Second Law of Thermodynamics tells us that the net process will only happen if the net entropy change is positive. However, the Second Law is more accurately viewed as a statistical regularity rather than a rigid law. The Fluctuation Theorem makes this explicit, relating the probabilities of a forward and reverse change during a given time interval to the associated entropy change. Although only proven recently (6), the basic idea has been an underlying theme in earlier works (such as (7)). Consider a simple transformation; for example, the forward direction could be the transformation of 1 ATP molecule to 1 ADP molecule plus 1 P_i_, and the reverse is the synthesis of 1 ATP from 1 ADP. During a time interval *t*, we call P_+_ the probability of the forward transformation happening and P_−_ the probability of the reverse transformation happening. If the entropy change associated with the forward transformation is ΔS, the Fluctuation Theorem relates the two probabilities as:

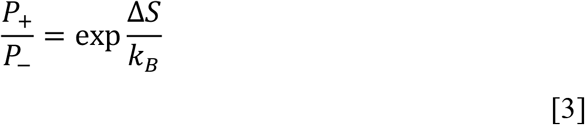

When ΔS is much larger than *k*_*B*_, the forward direction of change is overwhelmingly favored relative to the reverse direction of change, consistent with the Second Law.

Based on this relation, the rate of any transformation (e.g., diffusion down a gradient or a chemical reaction) can be expressed as (1):

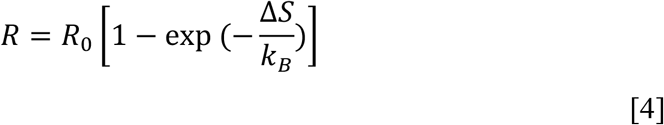

We can think of this as the product of two terms: a kinetic term (R_0_) and a thermodynamic term involving the entropy change. The kinetic rate R_0_ is set by mechanisms such as enzyme kinetics or diffusion, and is the rate the reaction will have if the entropy change (ΔS) is much larger than Boltzmann’s constant k_B_. As ΔS approaches zero, though, the rate of the reaction goes to zero due to the thermodynamic term.

### 2.3 Restoring ATP with oxidative metabolism

In the brain, ATP is consumed in cellular work and then restored primarily by coupling ATP synthesis to the oxidative metabolism of pyruvate in the mitochondria. Oxidative metabolism is a tightly coupled chain of reactions ending with the conversion of ADP to ATP (8). Along this path, intermediate products of one reaction in the chain are consumed by the next reaction in the chain (e.g., protons are pumped across the mitochondrial inner membrane and then return to drive ATP synthesis). By Eq [4], the net rate of metabolism is affected by the net entropy change of this chain of reactions, ΔS_net_. Ideally, the intermediate products along the chain, if they are produced and consumed in equal numbers, do not contribute to the net entropy change, and ΔS_net_ can be considered just in terms of the entropy changes of the first and last steps, as described by Eqs [1-2]. In terms of the numbers involved, restoring one molecule of ATP has a negative entropy change -ΔS_ATP_, and oxidative metabolism of one molecule of pyruvate has a positive entropy change ΔS_OX_. Empirically, the oxidative metabolism of 1 pyruvate is coupled to the conversion of 17 ADP to 17 ATP, so the net entropy change of the coupled reactions is then ΔS_net_ = ΔS_OX_ – 17ΔS_ATP_.

In principle, an efficient metabolism should generate the most ATP possible for each pyruvate molecule metabolized. This suggests that ΔS_net_ normally should be large enough that it does not limit the metabolic rate by Eq [4]. Increasing baseline ΔS_net_ further, though, would be wasteful in the sense that more nutrients (glucose and O_2_) are required to fuel the same level of ATP synthesis. However, this efficiency means that the metabolic rate would be more sensitive to a reduction of ΔS_net_ due to a reduction of tissue O_2_/CO_2_.

### 2.4 The distinction between maintaining the O_2_ metabolic rate and maintaining the phosphorylation potential at low O_2_

We now turn to how the considerations of entropy described in Eqs [1-4] might be expected to play out when tissue O_2_/CO_2_ is reduced. For a small reduction of ΔS_OX_, the net entropy change, ΔS_net_, will be reduced, but by Eq [4] this may have little effect on the overall metabolic rate. For a larger reduction of tissue O_2_/CO_2_, with a larger decrease of ΔS_net_, the rate of ATP production will be reduced. If the rate of ATP consumption in cellular work continues, though, the ratio of ATP/ADP will fall because the ATP is not being replenished. As this ratio falls, ΔS_ATP_ also will decrease (i.e., the phosphorylation potential will fall). If ΔS_ATP_ falls sufficiently, restoring ΔS_net_ to its normal value despite the fall in ΔS_OX_, the net metabolic rate would be restored. The overall effect is that the metabolic rate can be unaffected by the reduction of ΔS_OX_, but the major consequence is a parallel reduction of the phosphorylation potential. Because nearly all cellular work depends on harnessing the ΔS_ATP_ available from ATP, this corresponds to a major degradation of energy metabolism in the brain, despite the maintenance of the metabolic rate of ATP production.

This scenario, of a reduced phosphorylation potential despite a normal metabolic rate, is consistent with the important early work of Wilson and colleagues (9). Previous experiments had shown that the O_2_ metabolic rate is not limited by low PO_2_ until very low values are reached (PO_2_ < 1 Torr) (10), and the accepted view was that energy metabolism in general is not adversely affected until this very low level of PO_2_ is reached. Through their initial work, though, as well as later additional studies, Wilson and colleagues (11) found that the phosphorylation potential began to degrade when PO_2_ was reduced below about 12 Torr. In our earlier work using a simple model of O_2_ and CO_2_ transport (1), we re-analyzed previously reported *in vivo* brain hypoxia data from (12) that included measurements of the phosphorylation potential. The result was that with increasing hypoxia, the phosphorylation potential began to decrease when the modeled tissue PO_2_ was reduced below Wilson’s critical 12 Torr (1). Experimental measurements of tissue PO_2_ in the brain (13, 14) indicate that it is normally about 25 Torr, providing a buffer (‘safety margin’) of only about a factor of two above the critical level where the phosphorylation potential begins to be impaired.

In short, the entropy considerations suggest a somewhat counterintuitive scenario. As PO_2_ decreases, even though the O_2_ metabolic rate can be maintained, the phosphorylation potential can degrade. Our focus here on the tissue O_2_/CO_2_ ratio as suggested by the entropy argument, rather than just the O_2_ level, brings in the possibility that manipulating the CO_2_ level can help to preserve tissue O_2_/CO_2_ and the entropy available from oxidative metabolism. This is an important factor in the modeling results on hypoxia and hypercapnia reported here. The implication of the O_2_/CO_2_ hypothesis is that effectively coping with hypoxia would be expected to involve physiological responses that serve to help maintain tissue O_2_/CO_2_ above that critical level of a ∼50% reduction from baseline, where the phosphorylation potential begins to degrade.

## 3. Methods: Modeling changes in tissue O_2_/CO_2_

### 3.1 Parameters, units, and basic relationships

The key parameters, with units and typical values, are described here. Because the modeling involves several interacting physiological variables, it is convenient to express them in compatible units; here we express concentrations in millimoles/L (mM), partial pressures of O_2_ and CO_2_ in Torr (mmHg), H^+^ concentration in standard pH units, and time in minutes (min).

#### Flow and O_2_ metabolism

Blood flow (CBF) is abbreviated as *F*, and defined as the volume of arterial blood delivered to the capillary bed in a tissue element per minute divided by the total volume of the tissue element. It is useful to think of *F* as essentially having units of min^-1^, with a typical human CBF of 50 ml blood/100g-min being about *F*=0.5 min^-1^. The O_2_ metabolic rate (CMRO_2_) is abbreviated as *R*_*O2*_, and is usually expressed as moles/100g-min. Here again the units can be simplified by noting that CMRO_2_ essentially has dimensions of concentration/time (mM/min), with a typical value for human brain of about 1.8 mM/min. The net oxygen extraction fraction is dimensionless and abbreviated as *E*, and in the human brain at rest a typical value is *E*=0.4 (15). For comparisons with the rat brain, we assumed *F* and *R*_*O2*_ are typically three times higher in the rat for the awake reference baseline state (16).

#### O_2_ transport in blood

Oxygen concentration as dissolved gas is expressed as an O_2_ partial pressure (PO_2_), in units of Torr (or mmHg), with 7.5 Torr = 1 kPa. The solubility of O_2_ in plasma is α_O2_ = 0.00134 mM O_2_/Torr (equivalent to 0.003 ml O_2_/100L-mmHg; i.e, 1 mM O_2_ concentration = 2.24 ml O_2_/100 ml at standard temperature and pressure). Most of the O_2_, though, is bound to hemoglobin. In conventional units, fully saturated hemoglobin with a concentration of 15 g/100ml, with 1.34 ml of O_2_ bound per gram of hemoglobin when fully oxygenated, has an equivalent total O_2_ concentration of 20.1 ml/100ml. This is equivalent to an O_2_ concentration of about 9.0 mM, and we define the typical effective hemoglobin concentration as *H* = 9.0 mM (i.e., *H* is expressed in milliequivalents of O_2_, the total concentration of O_2_ that would result if the hemoglobin is fully saturated). The concentration of hemoglobin-bound O_2_ is then *S*_*O2*_*H*, where *S*_*O2*_ is the fractional saturation of hemoglobin binding sites for O_2_. The value of PO_2_ for which hemoglobin is 50% saturated (*S*_*O2*_ = 0.5) is designated P_50_, with a typical value in humans of 26.8 Torr, and for the rat we assumed 38 Torr (17-19).

#### CO_2_ transport in blood

The CO_2_ concentration as a dissolved gas is also expressed as a partial pressure, with a solubility α_CO2_ = 0.0307 mM/Torr. A larger component of CO_2_ in blood is in the form of bicarbonate ions (HCO_3_^-^), and we assume PCO_2_ and bicarbonate quickly approach equilibrium as PCO_2_ changes. In addition, a component of CO_2_ in blood is bound to hemoglobin, and the concentration of this component is *S*_*CO2*_*H*, where *S*_*CO2*_ is the fraction of CO_2_ binding sites bound to CO_2_.

#### H^+^ concentration

Finally, H^+^ concentration is expressed in normal pH units, with a typical arterial value of 7.4. In the modeling to follow we need to consider how pH changes down the length of a capillary in tissue, as it affects both the bicarbonate concentration, *S*_*CO2*_ and the value of P_50_. This relationship is complicated to model because O_2_ is being subtracted as CO_2_ is added to the blood. In our approach we assume standard buffering curves for fully oxygenated hemoglobin and fully deoxygenated hemoglobin with a linear variation between these curves as S_O2_ changes (described in the **Appendix**).

The standard values of physiological parameters assumed for a normal baseline state are summarized in **Table 1**. Values of parameters of the normal baseline state calculated with the model in the following section are listed in **Table 2**.

**Table 1.** Reference baseline state for the human brain: assumed parameters

| Parameter | Definition | Assumed Baseline Value |
| --- | --- | --- |
| $F$ | Cerebral blood flow | $0.5 \text{ min}^{-1}$ |
| $E$ | $\text{O}_2$ extraction fraction | 0.4 |
| $H$ | Arterial hemoglobin in milliequivalents of $\text{O}_2$ at 100% saturation | 9 mM<br>(15 g/dL) |
| $R_{\text{O}_2}$ | Cerebral metabolic rate of $\text{O}_2$ | 1.8 mM/min |
| $\text{pH}_a$ | Arterial pH | 7.4 |
| $\text{P}_a\text{O}_2$ | Arterial $\text{O}_2$ partial pressure | 90 Torr |
| $\text{P}_a\text{CO}_2$ | Arterial $\text{CO}_2$ partial pressure | 40 Torr |
| $\text{P}_{50}$ | $\text{PO}_2$ for half saturation of hemoglobin | 26.8 Torr |
| $\alpha_{\text{O}_2}$ | $\text{O}_2$ solubility | 0.00135 mM/Torr |
| $\alpha_{\text{CO}_2}$ | $\text{CO}_2$ solubility | 0.0307 mM/Torr |

**Table 2.** Reference baseline state for the human brain: calculated parameters based on the model.

| Parameter | Definition | Calculated Baseline Value |
| --- | --- | --- |
| $\text{S}_a\text{O}_2$ | Arterial $\text{O}_2$-hemoglobin saturation | 0.97 |
| $\text{T}_{\text{O}_2}$ | Total arterial $\text{O}_2$ | 8.84 mM |
| $\text{S}_a\text{CO}_2$ | Arterial $\text{CO}_2$-hemoglobin saturation | 0.056 |
| $\text{T}_{\text{CO}_2}$ | Total arterial $\text{CO}_2$ | 26.7 mM |
| $\text{P}_c\text{O}_2$ | Average capillary $\text{PO}_2$ | 47.9 Torr |
| $\text{P}_c\text{CO}_2$ | Average capillary $\text{PCO}_2$ | 46.0 Torr |
| $\text{P}_t\text{O}_2$ | Assumed average tissue $\text{PO}_2$ (baseline) | 25 Torr |
| $\tau$ | Tissue diffusion time constant (for assumed $\text{P}_t\text{O}_2=25$ Torr) | 0.017 min (1.02 s) |
| $\text{P}_t\text{CO}_2$ | Average tissue $\text{CO}_2$ | 47.0 Torr |
| $r$ | Tissue $\text{O}_2/\text{CO}_2$ (as $\text{PO}_2/\text{PCO}_2$) | 0.53 |

### 3.2. Basic equations and approach in the modeling

The rate of oxygen metabolism (*R*_*O2*_) can be expressed in two ways. The basic equation of mass balance for oxygen metabolism is:

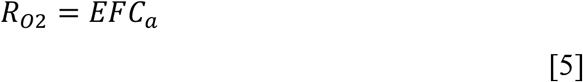

where *F* is the blood flow, *E* is the net O_2_ extraction fraction, and *C*_*a*_ is the arterial concentration of O_2_ (*R*_*O2*_ and *F* are a shorthand notation for CMRO_2_ and CBF, respectively; see **Table 1** for typical normal human values). The quantity *FC*_a_ is simply the rate of O_2_ delivery to a tissue element by blood flow, and *E* is then the fraction that is metabolized. For a steady-state, oxygen metabolism also can be considered as a diffusive flux of O_2_ (as a dissolved gas) down a gradient from the PO_2_ in the blood plasma to the PO_2_ in the tissue, modeled as:

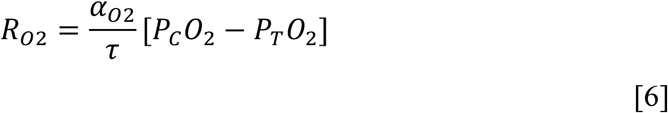

In this equation, *P*_C_O_2_ is the average PO_2_ in the capillary, *P*_T_O_2_ is the average PO_2_ in the tissue space, and the solubility α_O2_ scales these values to equivalent concentrations (i.e., the concentration in the fluid for a given partial pressure). The time constant τ depends on the diffusion coefficient of O_2_ and the capillary bed geometry.

Although this equation deals only with average values of PO_2_, its derivation does not require that PO_2_ is uniform in the tissue space. In the **Appendix Section A1**, this equation is motivated by the example of the Krogh cylinder model, which explicitly models spatial variability of PO_2_. In the context of that model, the time constant τ scales with the typical time required for an O_2_ molecule to diffuse to the farthest point in tissue from a capillary. For this reason, we might expect that τ will decrease for a capillary bed that occupies a greater fraction of the tissue volume (i.e., closer capillary spacing). However, we should take this only as a rough guide because the brain capillary bed is much more complex than the simple Krogh cylinder model (e.g., see the full capillary bed renderings in (20) and (21)).

A similar equation to Eq [6] applies to CO_2_ diffusion (described in the **Appendix Section A1**), with the major difference that the solubility of CO_2_ is about 23 times higher than that of O_2_. As a consequence, the difference between capillary and tissue PCO_2_ is about 23 times smaller than for the same difference in PO_2_. Here we have assumed that the diffusion constants for O_2_ and CO_2_ are the same, as discussed in the **Appendix Section A1**. Including a slight difference would alter the value of τ for CO_2_ by the ratio of the diffusion constants, a small effect compared to the large effect due to the ratio of solubilities. With this approximation, we can estimate tissue O_2_/CO_2_ as:

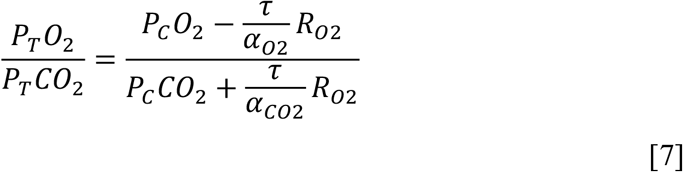

Here we express tissue O_2_/CO_2_ as a partial pressure ratio; the concentration difference can be calculated by multiplying by the respective solubilities.

To estimate tissue O_2_/CO_2_ with Eq [7] requires first an estimate of mean capillary PO_2_ and PCO_2_ under different conditions, and our approach for calculating these values is described in the **Appendix Sections A2-A6**.

The parameter τ is determined by the capillary bed geometry and the O_2_ diffusion properties, and captures the microscopic details of the diffusion process from capillary to mitochondria. This includes diffusion in tissue and any permeability limitation at the capillary wall. These microscopic details are not well known for the human brain. For this reason, in the calculations to follow we do not assume an *a priori* estimate of τ, but rather treat τ as a parameter that is determined from the baseline state (**Table 1**) by requiring consistency with Eqs [5-6]. To do this, we assume a value for the tissue PO_2_ for the baseline state, P_T_O_2_ = 25 Torr, a value consistent with experimental measurements (14, 22). After estimating average capillary PO_2_ and PCO_2_ using the model in **Appendix Sections A2-A6** (results listed in **Table 2**), the assumption of this value for tissue PO_2_ leads to an estimate of τ = 1.02 s for the human brain.

In applications of the model to estimate how tissue O_2_/CO_2_ changes in different physiological states, we assume this value of τ is unchanged. To evaluate a new physiological state, with potentially different values of CBF, CMRO_2_ and the arterial parameters, the first step is to estimate new values for the average capillary PO_2_ and PCO_2_, with the extraction fraction *E* determined by Eq [5]. From these average capillary values, the tissue O_2_/CO_2_ ratio is calculated as the ratio of partial pressures with Eq [7]. Our primary outcome variable is then the ratio *r* of tissue O_2_/CO_2_ in the new state compared to the reference baseline state:

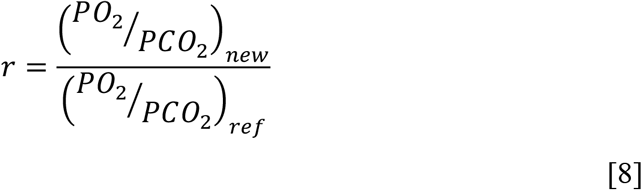

A reduction of 50% (*r*=0.5) is interpreted as the threshold for impairing the phosphorylation potential based on the analysis of hypoxia data in (1) and the earlier work of Wilson and colleagues (11).

## 4. Results

### Changing CMRO_2_ in normoxia

The primary way we illustrate the effects of the different variables is by plotting oxygen metabolism (CMRO_2_) on the *x*-axis and blood flow (CBF) on the *y*-axis. On each axis the values are normalized to the baseline state values. For the reference baseline state defined in **Table 1**, there is a curve in this plane corresponding to maintaining tissue O_2_/CO_2_ at the reference level as CMRO_2_ is changed (the blue curve in **Figure 1**). In this plot the dynamic range of CMRO_2_ is about 30%, corresponding to variations seen during normal brain function (e.g., during brain activation experiments). For example, the prediction of the modeling is that a CMRO_2_ increase of 20% requires a 60% increase of CBF to preserve tissue O_2_/CO_2_ at the baseline reference level. This much larger change in CBF creates the blood oxygenation level dependent (BOLD) effect exploited in functional MRI studies to map patterns of activation in the brain. In addition, the critical line showing the level at which the phosphorylation potential begins to degrade also can be plotted in this plane (the red curve in **Figure 1**). A CMRO_2_ increase of 30% with no change in CBF (indicated by the black square) would put the brain state near this critical level.

**Figure 1.**
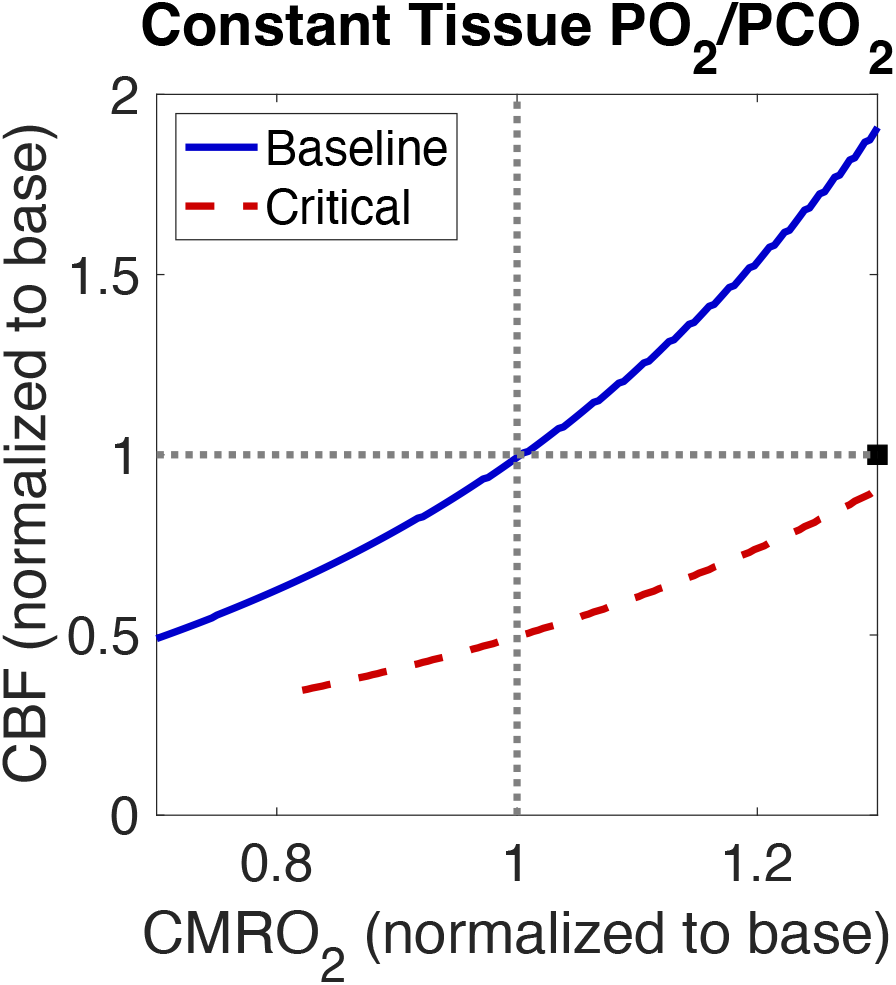
Theoretical curves for constant tissue O_2_/CO_2_. CBF and CMRO_2_ are normalized to their baseline values (**Table 1**). The curves show the CBF necessary to maintain baseline tissue O_2_/CO_2_ as CMRO_2_ varies around the baseline value (solid blue line) and the critical curve for avoiding degradation of the phosphorylation potential (dashed red line). A 30% increase of CMRO_2_ with no change in CBF (black square) is close to the critical state.

### Comparing human and rat brain

There are two critical quantitative differences for the rat compared to the human brain: CBF and CMRO_2_ are about three times higher in the awake baseline state of the rat (23), and the value of P_50_, the PO_2_ for half-saturation of hemoglobin, is larger (∼38 Torr compared to ∼27 Torr). Assuming these differences, with the rest of the reference baseline state in **Table 1** the same, we calculated the corresponding curves appropriate for the rat (**Figure 2**). To do this we adopted the same convention of estimating τ by assuming that the tissue PO_2_ is 25 Torr in the baseline state, which gives a value of 0.51 s for the rat, about half the value estimated for the human. As developed in **Appendix A1** we expect τ to be smaller when the capillary density is higher. A recent study (21), based on complete renderings of the capillary beds in sections from mouse and human brains, concluded that many of the topological properties of the two networks can be understood as an increased spatial scale for the human brain, with capillaries farther apart and longer. This experimental result, although for the mouse, is consistent with our estimate of a lower value of τ for the rat.

**Figure 2.**
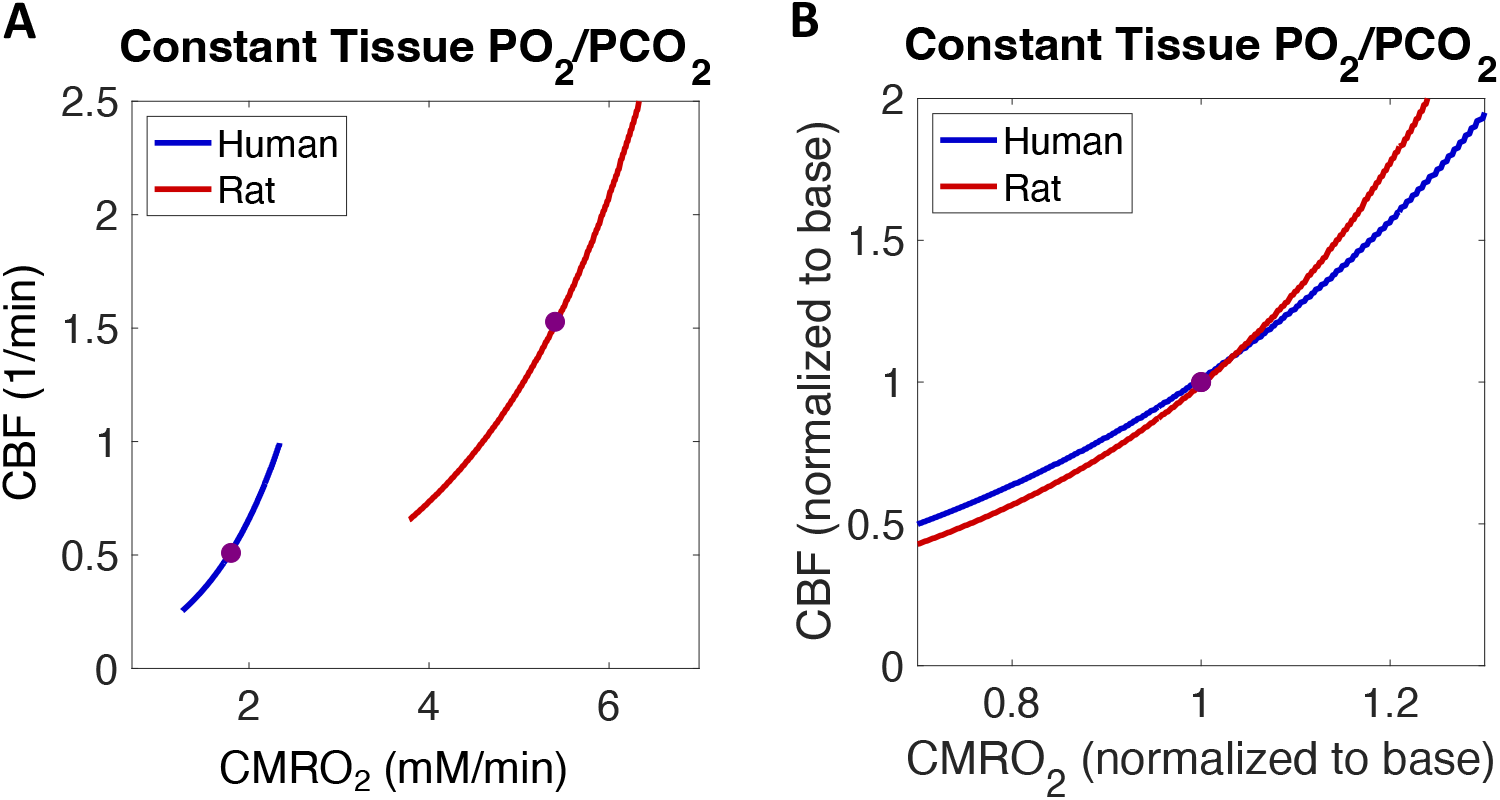
Comparing the human and rat for maintaining the same baseline tissue O_2_/CO_2_. **A**) Curves are plotted in the CBF/CMRO_2_ plane in absolute units, illustrating the higher values in the rat. The black circles indicate the awake resting baseline state. **B**) When normalized to their respective baseline values the curves are similar.

### Similar relationships for rat and human brain when normalized to baseline values

In **Figure 2A** the CBF and CMRO_2_ values are plotted as absolute values to show the wide spread between the rat and human values. However, when each curve is plotted normalized to the respective baseline state (**Figure 2B**), the curves are similar. **Figure 3** shows a number of experimental results in humans and rats, compared with the theoretical curve for humans. The human brain activation results are taken from both fMRI and PET studies, as summarized in (4). The rat data are from the review by Hyder and colleagues (23) of measurements for reduced CMRO_2_ (e.g., different stages of anesthesia). The theoretical curve for preserving tissue O_2_/CO_2_ is in good agreement with the experimental data.

**Figure 3.**
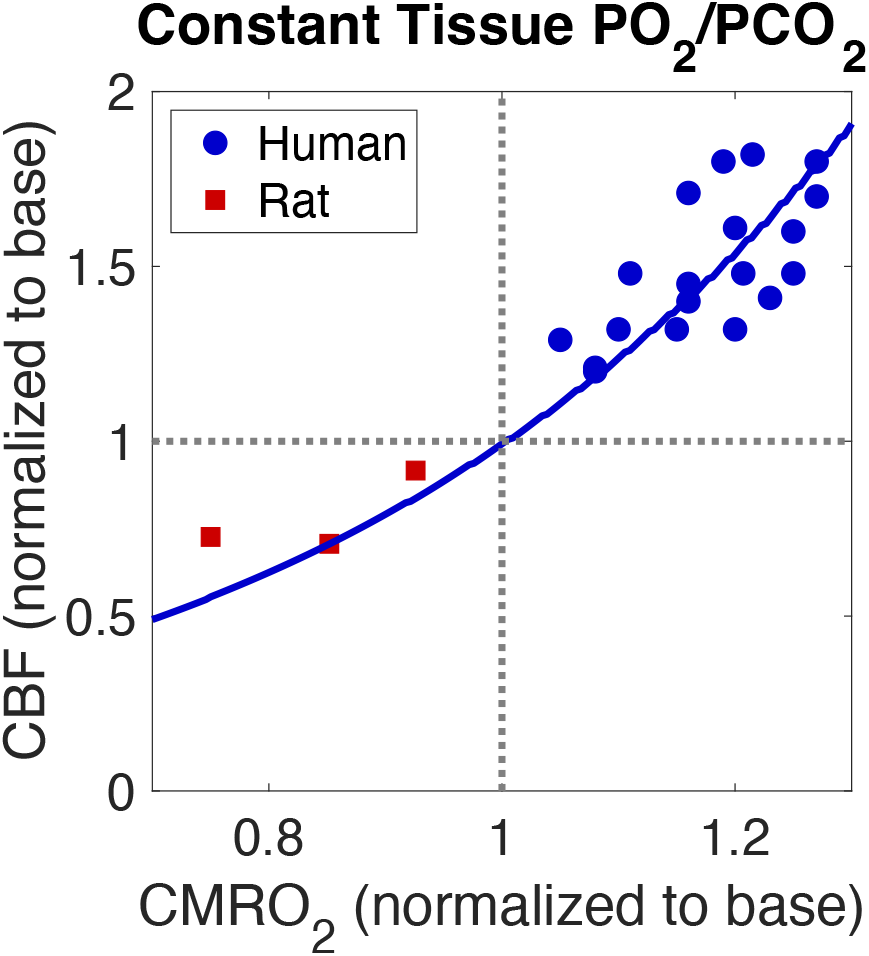
Theoretical curve for maintaining baseline tissue O_2_/CO_2_ compared with experimental data in humans and rats. The human data include the fMRI and PET studies listed in (4). The rat data are taken from (23).

### The role of metabolic scope

The brain differs from the heart in terms of basic features of O_2_ metabolism, and at first glance the combination of differences seems odd. Although baseline CMRO_2_ is high, the variation during normal awake brain function (metabolic scope) is low (∼30%), compared to the several fold changes in heart muscle. In addition, the baseline O_2_ extraction in the brain is lower, ∼40%, while for the heart it is 70-80% (24, 25). Intuitively, we might have expected that a lower extraction at baseline provides more capacity for accommodating a larger metabolic scope by simply increasing the extraction fraction. However, an important implication of the tissue O_2_/CO_2_ hypothesis is that there is a somewhat counterintuitive connection between baseline O_2_ extraction fraction, the value of τ (which depends on capillary density), and metabolic scope. To illustrate this connection, **Figure 4** compares the curves for preserving the same reference baseline value of tissue O_2_/CO_2_ for baseline O_2_ extraction fractions of 40% and 80%. For the same O_2_ metabolic rate the baseline CBF is reduced for the higher extraction fraction. To match the same tissue O_2_/CO_2_ ratio with the higher extraction fraction the value of τ also must be reduced, in this case to τ=0.52 s. Consistent with this implication of a shorter τ, the fractional blood volume in human brain is usually found to be about ∼4% (26), while that in the myocardium has been measured to be >8% (27). The interesting result is that for the higher baseline extraction fraction a much larger range of O_2_ metabolism can be accommodated with a given range of blood flow. The reason that the intuitive notion mentioned above is wrong is a key implication of the O_2_/CO_2_ hypothesis: to maintain tissue O_2_/CO_2_ the O_2_ extraction fraction must decrease with increased O_2_ metabolism. For this reason, increasing metabolic scope requires a smaller value of τ, and thus a higher capillary density, so that the baseline O_2_ extraction fraction can be higher.

**Figure 4.**
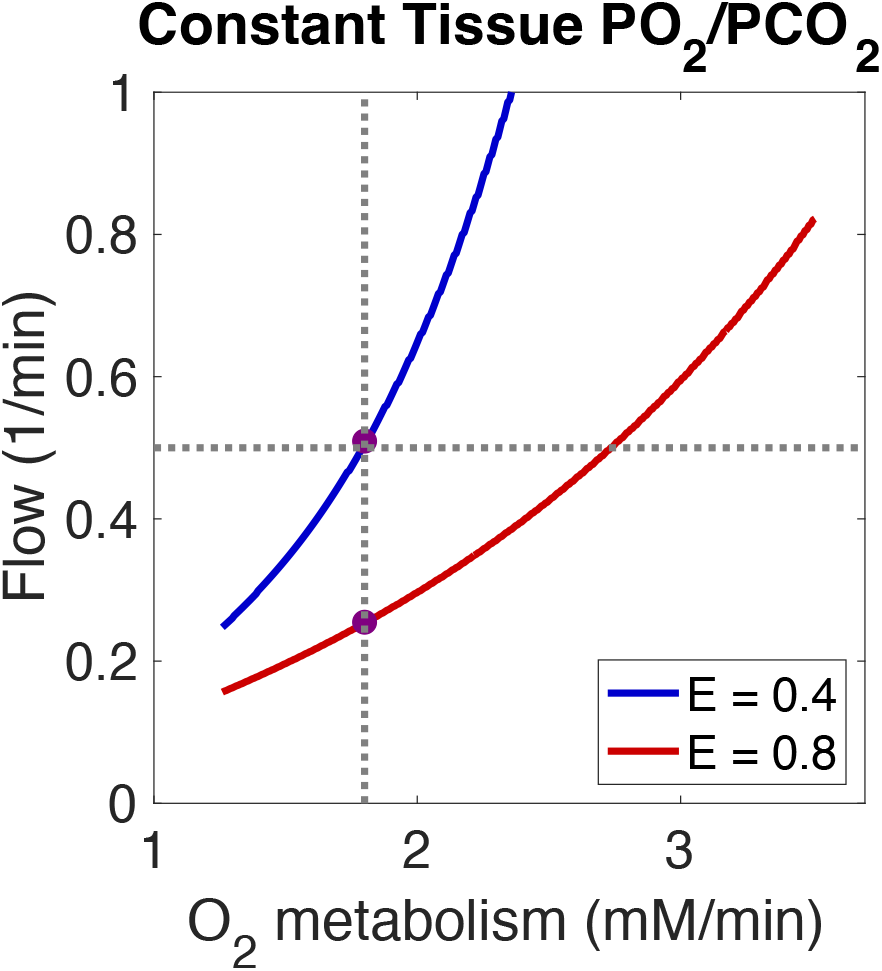
Baseline O_2_ extraction fraction, the value of τ, and metabolic scope. The curves for maintaining the same value of tissue O_2_/CO_2_ are compared for baseline O_2_ extraction fractions of 40% (typical for the brain) and 80%. For the same tissue O_2_/CO_2_ ratio, τ=1.02 s for 40% extraction and 0.52s for 80% extraction. Dotted lines show the baseline values for the brain. The same range of flow variation supports a larger range of O_2_ metabolism (metabolic scope) when E=0.8 compared to E=0.4.

### Changing CBF in response to hypercapnia

CBF is known to increase strongly when breathing a gas with increased CO_2_ content. This reactivity is usually quantified as the % change in CBF per Torr increase of PCO_2_, and in their review Ainslee and Duffin (28) found an average among several studies of ∼3.8 %/Torr over the PCO_2_ range from 35-55 Torr. We used the current model to test how much the CBF would need to increase to restore baseline tissue O_2_/CO_2_ if arterial PCO_2_ is increased from the baseline value of 40 Torr to 50 Torr. For any change in the arterial state, such as hypercapnia, we can plot how the curve for preserving the baseline tissue O_2_/CO_2_ is shifted in the CMRO_2_/CBF plane (**Figure 5)**. To restore tissue O_2_/CO_2_, CBF needs to increase by about 40% if the baseline CMRO_2_ stays constant. This corresponds to a CO_2_ reactivity of 4%/Torr, in good agreement with the experimental data. In order to support an increase of CMRO_2_ in the hypercapnic state the CBF needs to increase further. This example illustrates the importance of thinking of CBF as a way to modulate tissue PO_2_, rather than just delivering O_2_ to the capillary bed. That is, in the context of the tissue O_2_/CO_2_ hypothesis, simply maintaining O_2_ delivery is not enough. Even though the CBF response to hypercapnia for baseline CMRO_2_ has already increased O_2_ delivery substantially, CBF needs to increase still further to accommodate an increase of CMRO_2_. This is in good agreement with an experimental study of brain activation during hypercapnia in humans (29).

**Figure 5.**
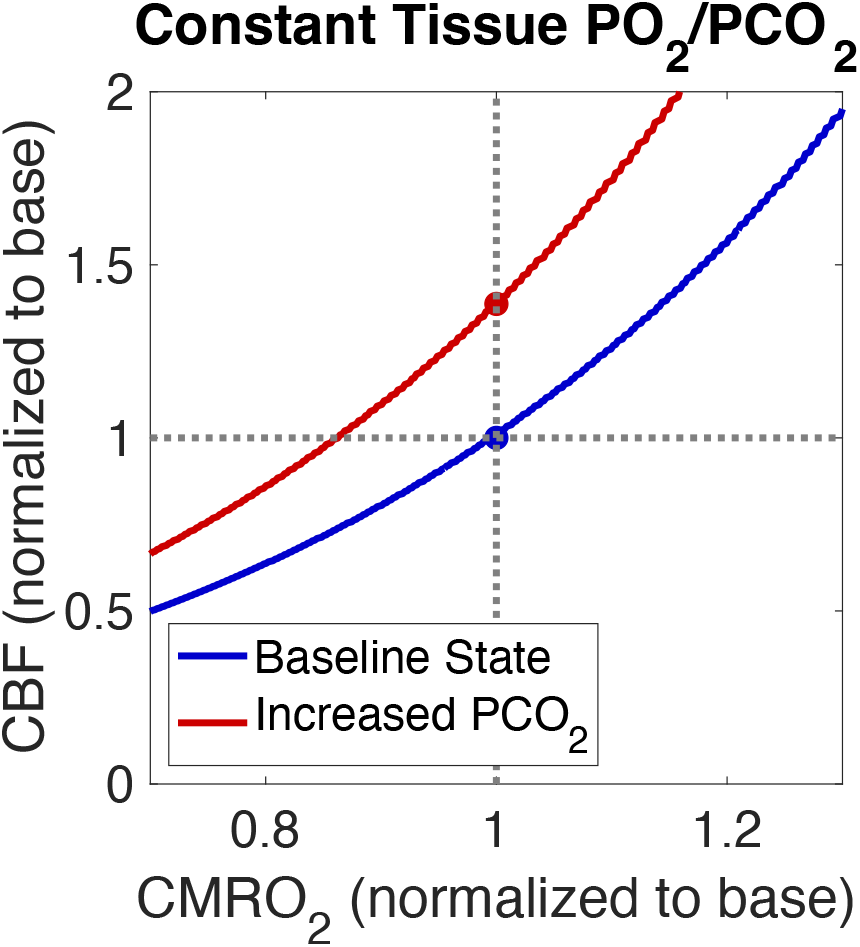
Effect of hypercapnia on maintaining tissue O_2_/CO_2_. The blue curve is the same as in **Figure 3**, showing the CBF change needed to maintain baseline tissue O_2_/CO_2_ for PCO_2_ = 40 Torr. If PCO_2_ is raised to 50 Torr the corresponding curve (red line) for maintaining the same value of tissue O_2_/CO_2_ is raised because the arterial PCO_2_ increase raises tissue PCO_2_. To restore the baseline tissue O_2_/CO_2_ value, CBF needs to increase by ∼40%. Although O_2_ delivery is already increased in the baseline state with elevated CO_2_, to accommodate an increase from baseline of CMRO_2_ requires a further increase in CBF.

### Compensating for hypoxia due to reduced arterial PO_2_

**Figure 6** shows how the curve for preserving tissue O_2_/CO_2_ shifts when arterial PO_2_ is lowered. **Figure 6A** shows the shift for a pure drop in arterial PO_2_ to 60 Torr (i.e., the other parameters of the baseline state remain the same). The curve is dramatically shifted up, and to restore tissue O_2_/CO_2_ requires CBF to increase by ∼90%. **Figure 6B** shows the effect of maintaining baseline O_2_ delivery by increasing the hemoglobin concentration to compensate for the reduced O_2_-saturation. Although this lowers the curve somewhat, the CBF change required to restore tissue O_2_/CO_2_ is still ∼77%. In contrast, **Figure 6C** shows that reducing arterial CO_2_ is an effective way to restore tissue O_2_/CO_2_ in hypoxia. In this case, a reduction of arterial PCO_2_ from 40 Torr to 28 Torr provides nearly full recovery, with only a small CBF increase of 5% needed to preserve tissue O_2_/CO_2_. In short, in the context of the tissue O_2_/CO_2_ hypothesis, reducing PCO_2_ is a critical factor for coping with hypoxia. Empirically, for humans acutely exposed to a reduction in PO_2_, hypoxic ventilatory stimulation results in a concomitant reduction in PCO_2_.

**Figure 6.**
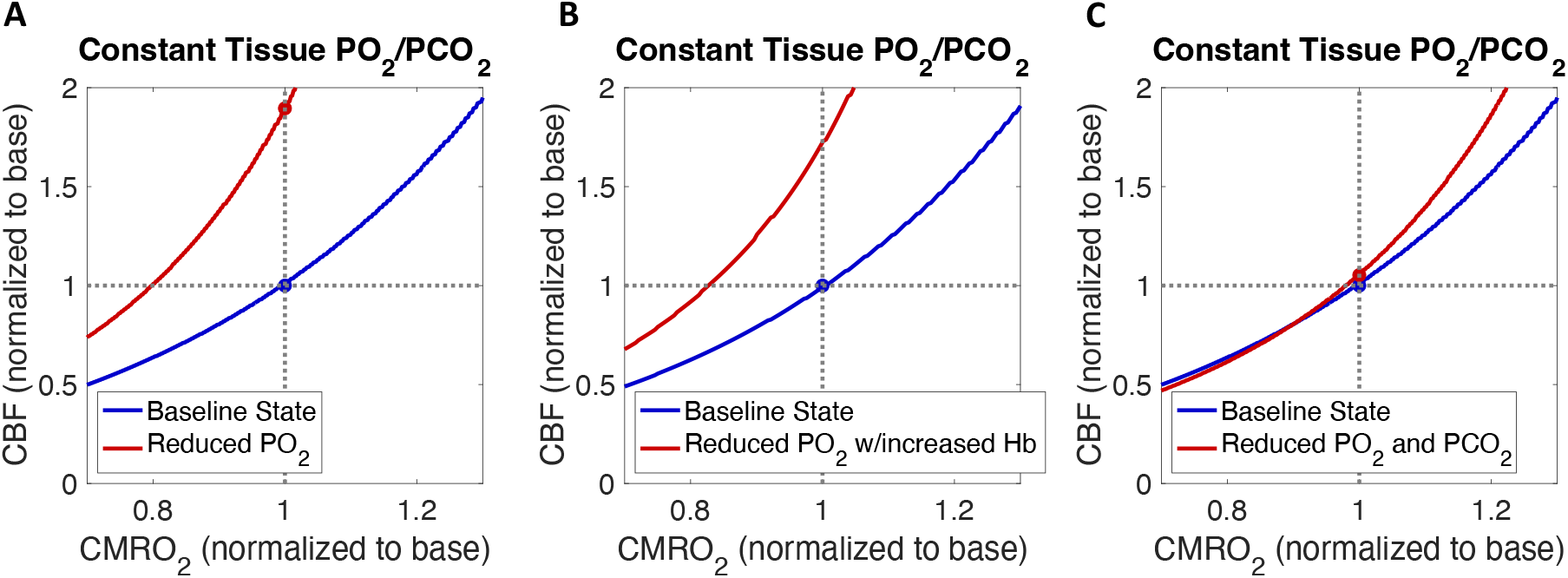
Effect of hypoxia on maintaining tissue O_2_/CO_2_. **A**) For a pure drop in PO_2_ to 60 Torr without a drop in PCO_2_ the curve for baseline tissue O_2_/CO_2_ is strongly shifted up, and a CBF increase of ∼90% would be required to restore the baseline level. **B**) For the same PO_2_ reduction, but now combined with increased hemoglobin concentration sufficient to maintain O_2_ delivery at the baseline level, the curve is still strongly shifted up and would require a CBF increase of ∼77% to restore baseline tissue O_2_/CO_2_. **C**) In contrast, combining the PO_2_ reduction with a concomitant reduction of PCO_2_ to 28 Torr essentially maintains baseline tissue O_2_/CO_2_ for the baseline value of CBF.

### Is tissue O_2_/CO_2_ preserved with acclimatization to high altitude?

When a healthy subject living at sea level travels to high altitude they exhibit a range of physiological changes as they acclimatize to the lower atmospheric PO_2_. **Table 3** lists the mean arterial state for subjects living at sea level who spent 3 weeks at an altitude of 5,260 m, from the study of Moller and colleagues (30). In this study baseline CBF and CMRO_2_ were also measured and found to be unchanged from sea level. **Figure 7** shows how the curve for maintaining the reference baseline tissue O_2_/CO_2_ is shifted for the altered arterial state. Interestingly, the acclimatized lowlanders precisely match the baseline tissue O_2_/CO_2_ for the normal values of CBF and CMRO_2_, in good agreement with the experimental findings that these were unchanged. However, the modeling predicts that for normal increases in CMRO_2_ with increasing neural activity there is a much steeper requirement for increased CBF at altitude compared to sea level. For example, supporting a 20% increase of CMRO_2_ at sea level requires a 54% increase of CBF, while for the acclimatized group a CBF change of 100% is required.

**Table 3:** Arterial state after acclimatization to high altitude.

|  | Lowlanders (Acclimatized for 3 weeks)<sup>A</sup> |
| --- | --- |
| Elevation (m) | 5,260 |
| PO<sub>2</sub> (Torr) | 51 |
| PCO<sub>2</sub> (Torr) | 25 |
| SO<sub>2</sub> (%) | 84 |
| Hemoglobin concentration (g/dL) | 17.9 |
| pH | 7.47 |
| P<sub>50</sub> (Torr)* | 28.2 |

A – from Moller et al (30)
\* – estimated with the model from SO<sub>2</sub> and PO<sub>2</sub> using Eqs [A6-A7]

**Figure 7.**
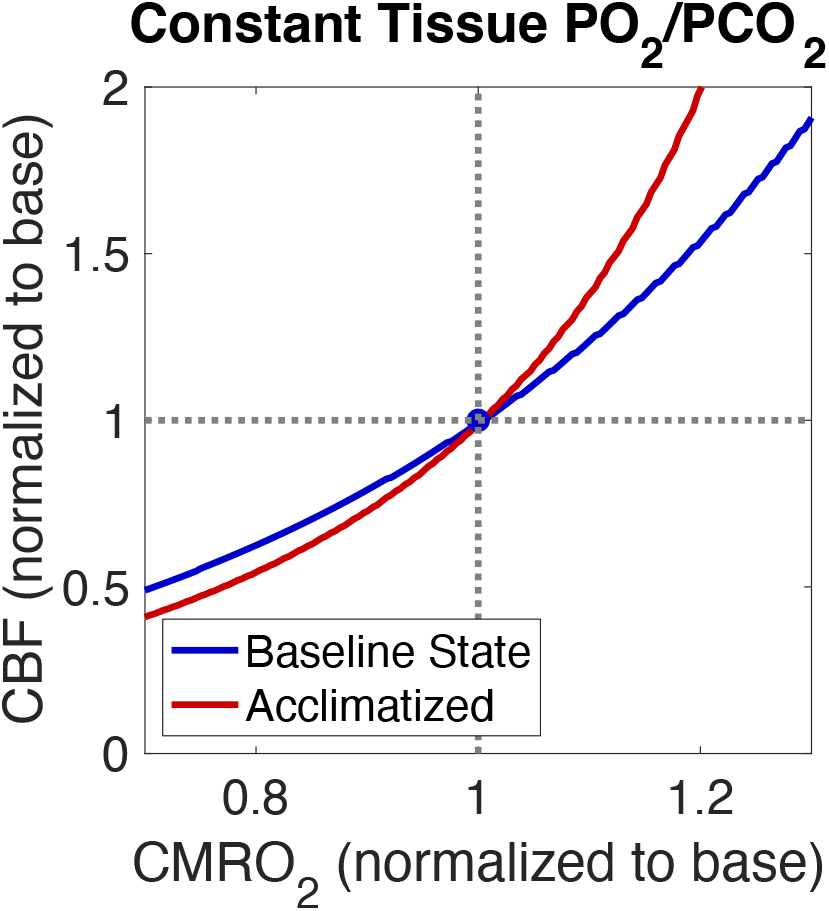
Effect of acclimatization to high altitude on tissue O_2_/CO_2_. The standard reference curve for maintaining baseline tissue O_2_/CO_2_ at sea level (with the arterial state in **Table 1**) is plotted in blue. For lowlanders acclimatized for 3 weeks to an elevation of 5,260 m (with the new arterial state in **Table 3**), the curve for maintaining tissue O_2_/CO_2_ is plotted in blue. The curves cross at the baseline CBF and CMRO_2_ values, consistent with the experimental finding that there was no change in these values between sea level and high altitude. However, in the acclimatized group a larger CBF change would be needed to support an increase of CMRO_2_ during normal brain function.

### How well do human populations adapted to high altitude preserve tissue O_2_/CO_2_?

Highland populations living in Tibet and the Andes at ∼4,200 m have adapted to cope with the low O_2_ environment. Estimates are that the Tibetan population has had about 25,000 years to adapt, while the Andean population has had about 11,000 years to adapt (31). Interestingly, the physiological strategies for these highlanders are somewhat different, with higher hemoglobin in the Andean population but lower PCO_2_ in the Tibetan population (31). Here we consider data reported for these populations and use the modeling to test how well tissue O_2_/CO_2_ in the brain is being maintained. **Table 4** shows the arterial state measured for the two populations, compiled from several sources given in the table. The assumption of an unchanged pH is based on a number of studies, summarized in Table 9.1 of (32). Taking these as the new arterial states, **Figure 8** shows the corresponding curves in the CMRO_2_/CBF plane required to maintain tissue O_2_/CO_2_. The curves suggest that tissue O_2_/CO_2_ has fallen, although not to a critical level, for the reference baseline values of CBF and CMRO_2_. However, it should be kept in mind that these estimates assume an unchanged value of τ from the reference state. If capillary density increases with adaptation to high altitude, reducing τ, these calculations will underestimate tissue O_2_/CO_2_. Although we are not aware of any brain capillary density measurements in these populations, studies in rats have found increased capillary density in response to chronic hypoxia (25). **Figure 9** shows the curves for maintaining the same tissue O_2_/CO_2_ if capillary density increases such that τ=0.88 s in both populations. This adaptation restores tissue O_2_/CO_2_ for the baseline CBF and CMRO_2_ values. However, as for the acclimatization response in **Figure 7**, the CBF change required to accommodate the normal range of CMRO_2_ variation in brain function is larger than for the baseline reference state. For example, a 20% increase of CMRO_2_ requires a CBF change of 54% in the baseline state. The needed increased CBF for the same CMRO_2_ change is greatest for the acclimatized lowlanders (∼100%), less so for the Andean population (∼77%), and still less for the Tibetan population (∼66%).

**Table 4:** Arterial state for populations adapted to high altitude.

|  | Andean Plateau<br>Natives | Tibetan Plateau<br>Natives |
| --- | --- | --- |
| Elevation (m) | ~4,200 | ~4,200 |
| PO<sub>2</sub> (Torr) | 53.1<sup>B</sup> | 55.9<sup>B</sup> |
| PCO<sub>2</sub> (Torr) | 32.3<sup>B</sup> | 29.6<sup>B</sup> |
| SO<sub>2</sub> (%) | 86.0<sup>B</sup> | 87.8<sup>B</sup> |
| Hemoglobin<br>concentration<br>(g/dL) | 19.2 (M)<sup>C</sup><br>17.8 (F)<sup>C</sup><br>18.5 (av) | 15.6 (M)<sup>C</sup><br>14.2 (F)<sup>C</sup><br>14.9 (av) |
| pH | 7.4 (assumed) | 7.4 (assumed) |
| P<sub>50</sub> (Torr)* | 27.8 | 26.8 |

B – from Moore (33)
C – from Beall (31)
\* – estimated with the model from SO<sub>2</sub> and PO<sub>2</sub> using Eqs [A6-A7]

**Figure 8.**
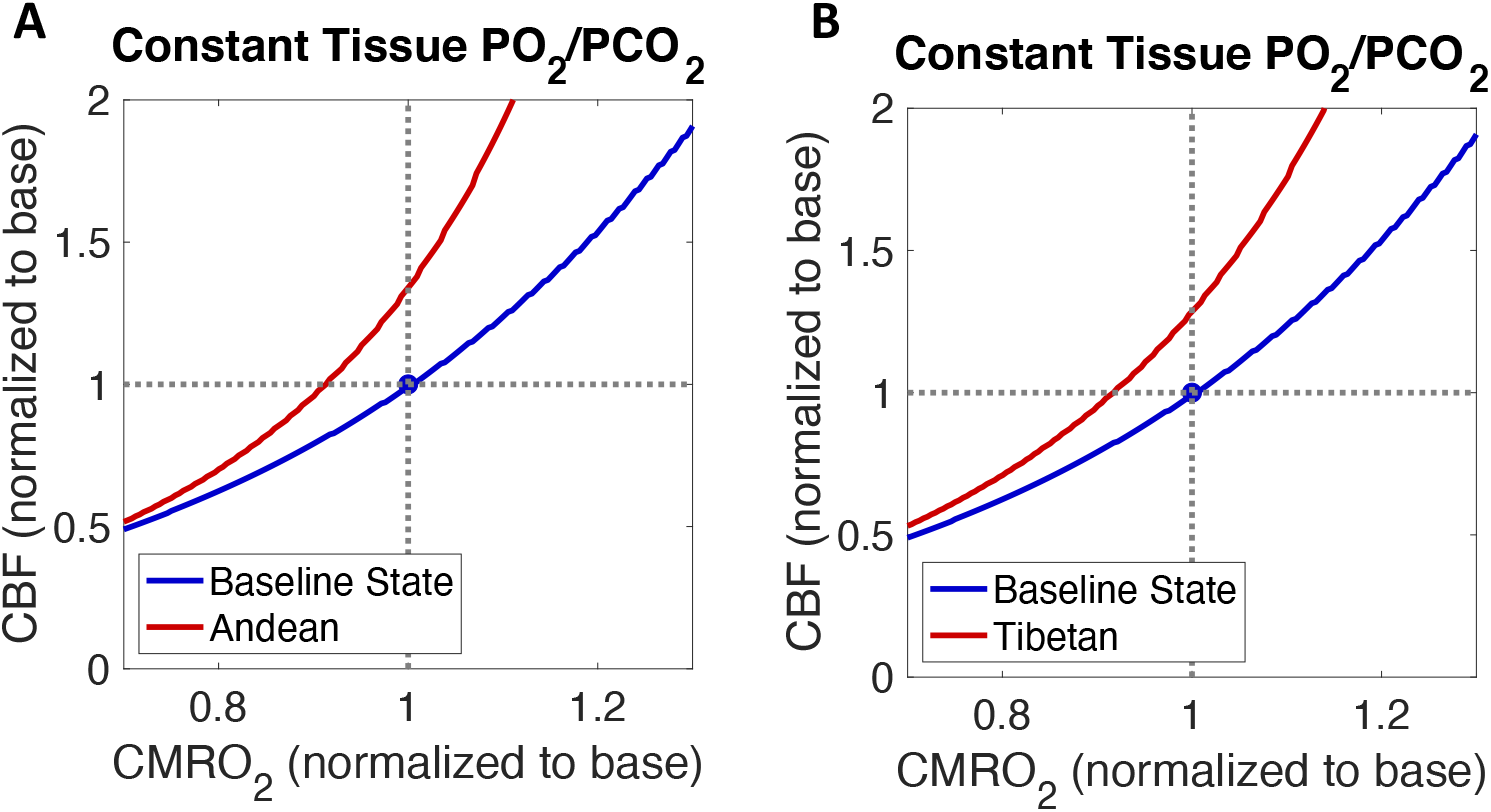
Estimates for maintaining tissue O_2_/CO_2_ for two populations adapted to high altitude. For the arterial states given in **Table 4**, the curve for maintaining the tissue O_2_/CO_2_ defined for the reference baseline state is plotted in the CMRO_2_/CBF plane. **A**) Andean plateau natives, and **B**) Tibetan plateau natives.

**Figure 9.**
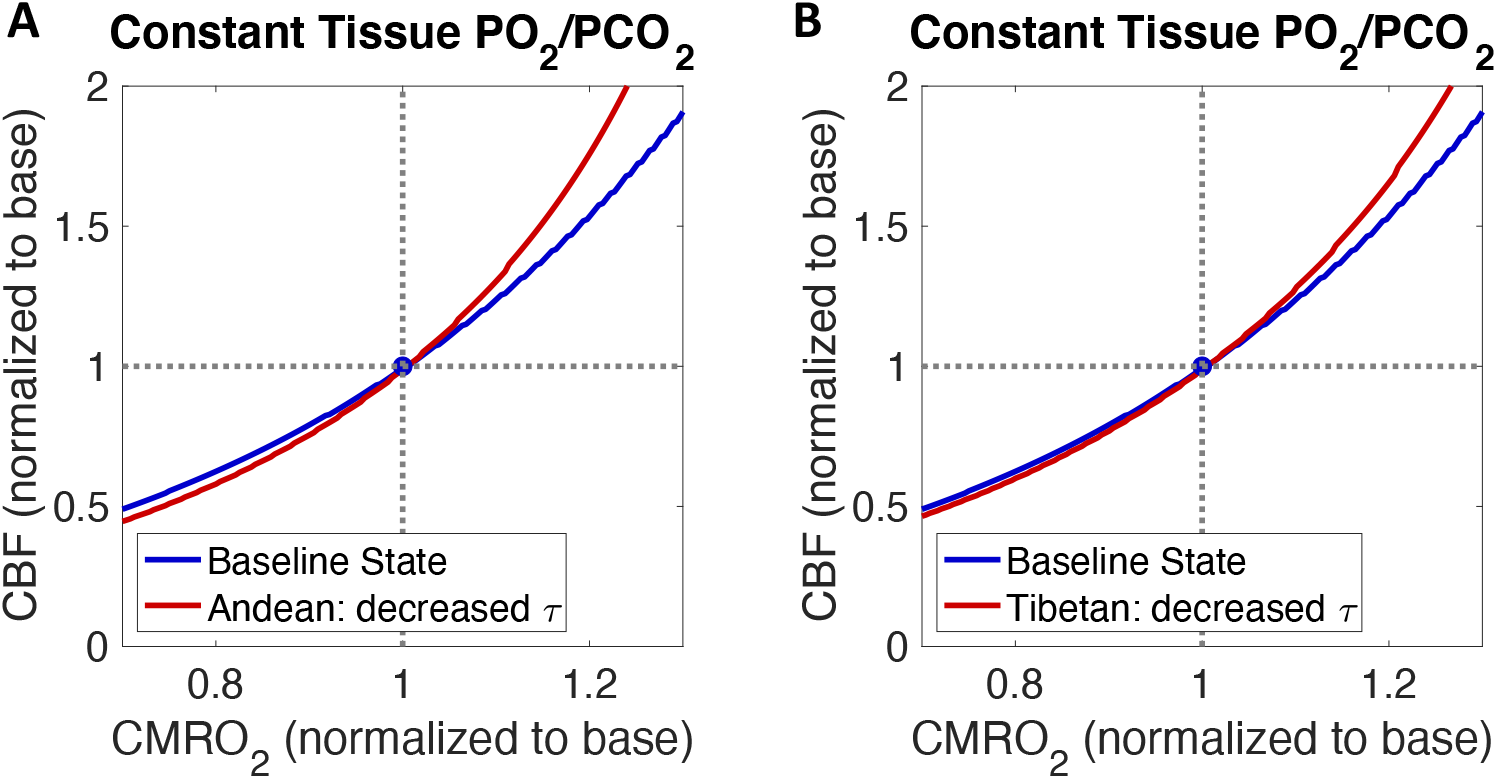
Possible role of increased capillary density in highland populations. Similar curves to those in **Figure 8**, with τ reduced from 1.02 s to 0.88 s (representing the effect of increased capillary density). **A**) Andean plateau natives, and **B**) Tibetan plateau natives.

## 5. Discussion

The central hypothesis tested here is that the physiological responses in the brain to a wide range of conditions act to preserve the ratio of O_2_/CO_2_ in tissue. In the initial paper describing the thermodynamic arguments for the hypothesis, a simplified model of O_2_ and CO_2_ transport was used to demonstrate the basic qualitative consistency of the hypothesis with experimental observations. Here that work is extended in two ways: 1) a more detailed model of O_2_ and CO_2_ transport was used, enabling more accurate predictions of the hypothesis; and 2) the model was then used to investigate the implications of the hypothesis for hypoxia in the brain. With the model, tissue O_2_/CO_2_ can be estimated given new values of CBF, CMRO_2_ and the arterial state, and compared with the value in a standard reference state. The hypothesis that the physiological changes act to preserve tissue O_2_/CO_2_ at the baseline level leads to predictions in good agreement with experimental results for:

- The large CBF increase associated with a smaller CMRO_2_ increase due to increased neural activity, the physiological basis of fMRI used to study brain activation based on the blood oxygenation level dependent (BOLD) effect.
- The large CBF increase associated with hypercapnia, a strong response that has been recognized for more than a century.
- The lack of a CBF increase in acclimatization to the low PO_2_ of high altitude when ventilation increases to lower PCO_2_. In this case no CBF change is needed to restore tissue O_2_/CO_2_ if arterial CO_2_ is sufficiently reduced when arterial PO_2_ is reduced.

The following sections discuss how the modeling results inform how we think about CBF and CMRO_2_.

### Function and mechanisms

This work focused on the function served by CBF, rather than the mechanisms that control CBF. The argument here is that preserving tissue O_2_/CO_2_ is important to the organism for preserving the phosphorylation potential. Interestingly, though, it does not appear that one cellular mechanism is responsible for accomplishing this function. The molecular mechanism that would be most useful to maintain tissue O_2_/CO_2_ would be one that is directly sensitive to that ratio, but there is currently no known sensor that functions in this way. In fact, many mechanisms have been identified that modify CBF (34-50), including direct effects of neural activity (important for the neural activation response) and pH (important for the response to hypercapnia). However, how these mechanisms fit together in an overall response is not well understood. For the response to increased neural activity, it appears that the large CBF changes are driven in a feed-forward way by aspects of neural activity, essentially increasing CBF in anticipation of an increase in CMRO_2_. Our analysis in terms of tissue O_2_/CO_2_ essentially defines two limits on how large such a CBF change needs to be: 1) the CBF required to maintain the normal baseline level of tissue O_2_/PO_2_ (i.e., maintaining the normal safety margin); and 2) the CBF required to maintain the phosphorylation potential, by keeping tissue O_2_/CO_2_ above a level of ∼50% of the normal baseline level (essentially using all the safety margin).

### CBF modulates tissue O_2_

Blood flow is often thought about in terms of delivery of O_2_ to the capillary bed: O_2_ delivery is simply CBF times the arterial concentration of O_2_. Several of the scenarios described here suggest that this is too limited a view. Of course, there needs to be adequate delivery of O_2_, but as long as CBF is high enough the important role suggested here is that modulation of CBF modulates tissue O_2_. For example, in hypercapnia the elevated arterial PCO_2_ raises tissue CO_2_ and lowers the O_2_/CO_2_ ratio. Increased CBF can restore tissue O_2_/CO_2_ by raising tissue O_2_. This results in a large increase in O_2_ delivery, but that is not the main function of the CBF modulation.

### Ventilation modulates tissue CO_2_

Increased ventilation has two effects relevant to the current ideas. The first is the well-known effect due to the alveolar gas equation, that for a given level of inhaled PO_2_, reducing the alveolar level of CO_2_ serves to increase the arterial PO_2_ achieved by gas exchange. For the current hypothesis, the second key role of ventilation is to reduce arterial CO_2_ to help compensate for reduced arterial PO_2_. That is, for any reduced arterial PO_2_ achieved by the lung, tissue PO_2_ will be reduced and the tissue O_2_/CO_2_ ratio can be restored by also reducing arterial PCO_2_. This scenario of reducing PO_2_ and PCO_2_ in parallel is essentially complementary to the hypercapnia picture, where tissue O_2_ is increased to compensate for increased PCO_2_.

### Potential conflict between function and mechanisms

CBF increases with increased CO_2_, a beneficial effect for preserving tissue O_2_/CO_2_ in hypercapnia. However, in a complementary way CBF decreases when CO_2_ is reduced, although not as steeply. However, this effect is counterproductive when CO_2_ is reduced to preserve tissue O_2_/CO_2_ in hypoxia. Successful acclimatization, which requires days or weeks, requires this mechanism to be reset, so that CBF is not reduced. The mechanism for the CO_2_ dependence of CBF is not fully understood, but there is evidence for the involvement of pH and, for hypercapnia, also nitric oxide (28). Normalizing pH by removing bicarbonate may be a critical component of adjusting this mechanism, as it is thought to be for resetting the dependence of ventilation on PCO_2_ (32).

### The diffusion of O_2_ from capillary to tissue

A key parameter in modeling the diffusion of O_2_ (Eq [3]) is a time constant τ, which depends on capillary geometry, the diffusion constant for O_2_ in tissue, and any permeability limitations as O_2_ diffuses out of the capillary. Here we did not try to model these factors, and instead estimated τ by requiring consistency between the modeling of O_2_ and CO_2_ transport and an empirical estimate of tissue PO_2_ in the baseline state of 25 Torr. This estimate of τ depends on the assumed value of the baseline O_2_ extraction fraction (*E*), which determines the mean capillary PO_2_, the driving head for diffusion into tissue. The primary interpretation of the physical meaning of τ, assuming it is dominantly determined by diffusion of O_2_ in tissue, is that it reflects capillary density, decreasing as capillary density increases.

### Baseline extraction fraction and the dynamic range of O_2_ metabolism

A primary result of the modeling (following from Eq [3]) is that to prevent tissue PO_2_ from falling when CMRO_2_ increases the average capillary PO_2_ needs to increase, which requires that the O_2_ extraction fraction (*E*) must decrease from the baseline value. The baseline value of *E* thus plays an important role related to a basic question of dynamic range: for a given dynamic range of blood flow, what dynamic range of O_2_ metabolism can be supported? We used the modeling to compare two values of the baseline *E*: 40% (typical of the brain) and 80% (more typical of the heart). For these two scenarios the estimated value of τ is higher for brain and the requirement for increased blood flow to support O_2_ metabolism is more severe. In short, the conclusion is that a higher capillary density (with associated smaller τ) enables a higher baseline O_2_ extraction fraction and provides a larger dynamic range for O_2_ metabolism for a given range of blood flow. This is consistent with what has been observed for the brain and heart, with a lower dynamic range and lower baseline E for the brain and a higher baseline E and larger dynamic range of O_2_ metabolism in the heart.

### The role of P_50_, the PO_2_ for half saturation of hemoglobin with O_2_

The value of P_50_ varies through several mechanisms, and an important effect is the generally higher value for smaller animals. Here we compared the predictions for rat brain compared to human brain. There are two important differences between the two species: 1) the values of CBF and CMRO_2_ are about 3 times higher in rat; and 2) the P_50_ value is higher in rat, about ∼38 Torr compared to ∼27 Torr in humans. In the modeling, the estimated value of τ (i.e., consistent with a baseline tissue PO_2_ of 25 Torr) was calculated from the absolute CMRO_2_ value and the appropriate value of P_50_. The result was a smaller τ for the rat, consistent with a higher capillary density for the same O_2_ diffusion properties compared to the human brain. The modeling showed that when the values of CBF and CMRO_2_ are expressed normalized to their respective baseline values, the fractional CBF change required to support varying fractional changes in CMRO_2_ is similar for the two species.

### Animals adapted to high altitude

A consistent finding in animals adapted to the low atmospheric O_2_ of high altitude is that the P_50_ values are lower than related species living at lower elevations (51). In the previous section we focused on the role of P_50_ in unloading O_2_ in the capillary bed, where the key factor is the average capillary PO_2_ as the driving head for diffusion into the tissue. However, the P_50_ also affects O_2_ loading from the lung. For sea level atmospheric PO_2_, the difference between the human and the rat P_50_ values has little effect on the hemoglobin O_2_ saturation of arterial blood. At higher elevations O_2_ loading becomes more critical, favoring a lower P_50_. However, to maintain brain tissue O_2_/CO_2_ requires either a lower PCO_2_ or an increased brain capillary density. Fully exploring this question in light of existing experimental data on animals adapted to hypoxia would be an important application for testing the tissue O_2_/CO_2_ hypothesis.

### The role of variable capillary density

The effects of capillary density, reflected in the value of τ, were considered in the context of variations across species for the same organ or across organs for the same species in the previous two sections. In the modeling, comparing tissue O_2_/CO_2_ in a new state with that in the reference baseline state, we specifically assumed that τ was unchanged. In principle, though, capillary density could vary in two ways: capillary recruitment, the opening of previously unperfused capillaries as blood flow increases; and growth of new capillaries, either within an individual organism or as part of genetic adaptation of a population. In the brain, experimental studies indicate that capillary recruitment is not an important effect during acute changes in CBF (52, 53). This is in sharp contrast to skeletal muscle, where capillary recruitment is thought to be an important factor. For this reason, the results here for the brain do not translate directly to muscle; if τ becomes smaller as flow increases, there would not be the same requirement for a large blood flow increase. For the brain, although we assume there is no capillary recruitment, studies have found evidence for increased capillary density in rat and mouse brain following prolonged exposure to hypoxia (25, 54). It is also possible that adaptations in populations who have lived at high altitude for many generations have led to a higher brain capillary density for the population compared to populations living at sea level, but to our knowledge this has not been tested.

### Physiological strategies involved in human adaptation to living at high altitude

As an initial test of the possible role of a higher capillary density, we compared the reported arterial states for Andean plateau natives and Tibetan plateau natives. Both populations have lived at elevations around 4,200 m for many generations (the Tibetans longer than the Andeans). Applying our assumption of the same value of τ as we calculated for our defined reference baseline (i.e., a sea level baseline), we estimated that the tissue O_2_/CO_2_ is reduced by about a factor of 0.84 in both populations. This is still above the critical reduction of 0.5 for beginning to degrade the phosphorylation potential. However, given that a higher capillary density is possible, we tested how much the value of τ would need to be reduced so that the tissue O_2_/CO_2_ is the same as for the baseline sea level state, giving a new value of τ=0.88 s compared to the baseline value τ=1.02 s. Interestingly, though, even with this adaptation to restore tissue O_2_/CO_2_ for the baseline values of CBF and CMRO_2_, the resulting curve of CBF needed to support a change in CMRO_2_ was steeper.

### The need to support variable CMRO_2_ over the normal range of varying neural activity

In the modeling tests of coping with hypoxia we focused on how the tissue O_2_/CO_2_ is altered for the baseline values of CBF and CMRO_2_ when the arterial state is changed. However, given the strong CBF change required to support a modest CMRO_2_ change, it is important to consider the range of CMRO_2_ required for normal brain function and the associated dynamic range needed for CBF. This may be important for thinking about possible cognitive effects, particularly for demanding tasks normally associated with increased neural activity. A consistent finding in the modeling of hypoxia was that the CBF curve as a function of increasing CMRO_2_ is steeper than the sea level baseline curve. This was most extreme for the acclimatization study, and less extreme for the two adapted highland populations (after adjusting for possible higher capillary density). Of the latter two populations, the curve for Tibetans was less steep, and approached the sea level curve. For example, in the baseline sea level state an increase of CMRO_2_ of 20% requires a CBF increase of 54%. For the three high altitude groups, the CBF change required for the same CMRO_2_ change is 100% for the acclimatized group, 77% for the Andean group, and 66% for the Tibetan group. In short, hypoxia significantly affects the dynamic range of CBF required to maintain tissue O_2_/CO_2_ over the range of variation of CMRO_2_ for normal brain function.

### The potential role of reducing CMRO_2_

In all the modeling studies an implicit assumption was that the CMRO_2_ required is determined by normal levels of neural activity. That is, the basic question was: how well can the normal level of brain function be maintained? However, it is important to consider the question of the consequences of reduced CMRO_2_. In terms of preserving the phosphorylation potential, reducing CMRO_2_ is an effective way to preserve tissue O_2_/CO_2_. For example, from **Figure 1** if CBF was reduced by 50% (e.g., due to stroke), the tissue O_2_/CO_2_ would be reduced to the critical level if CMRO_2_ continues at the baseline level. However, a reduction of CMRO_2_ by 10% would move the tissue O_2_/CO_2_ above the critical level, and a reduction of CMRO_2_ by 30% would restore baseline tissue O_2_/CO_2_. In short, such a reduction could preserve the phosphorylation potential for the remaining cellular work and improve survival of the tissue until CBF is restored.

### How can CMRO_2_ be reduced?

Inhibition of neural activity is the most direct way to lower CMRO_2_, by reducing neural activity. For example, tapping the fingers of one hand reduces CMRO_2_ in the brain region corresponding to the opposite hand through cross-callosal inhibition, as demonstrated in (55). In this study the authors also found a larger fractional CBF reduction, about 2.3 times larger than the fractional CMRO_2_ reduction, and consistent with the theoretical curve in **Figure 3** for the tissue O_2_/CO_2_ hypothesis. In coping with hypoxia it is possible that a more widespread inhibition could prevent the highest demand for CMRO_2_ and ease the requirement for a large CBF change. For example, in the modeling of acclimatized lowlanders compared to the high altitude native populations, the curve of required CBF change as a function of CMRO_2_ change is steeper. To limit the largest required CBF changes we would expect neural inhibition in the acclimatized lowlanders to be more severe than the native highlanders, potentially with associated cognitive impairment. For hypoxia due to stroke, a mechanism for increasing neural inhibition in response to the hypoxic effects could be beneficial, as discussed in the previous section. An important research question is to understand the physiological mechanisms that could connect hypoxia to increased neural inhibition (56-58).

The following sections discuss particular features and limitations of the modeling in relation to previous work as well as future directions.

### Comparison of the current modeling with our earlier work

In the earlier report on the tissue O_2_/CO_2_ hypothesis (1) we used a simpler model for the transport of O_2_ and CO_2_ to illustrate the basic consistency with observations. Here the modeling is based on recent models for O_2_ and CO_2_ binding to hemoglobin (59), combined with more accurate modeling of capillary O_2_ and CO_2_. We also considered species differences, with associated differences in the P_50_ for O_2_-hemoglobin binding. Based on the modeling, the resulting predictions of the hypothesis are in good agreement with observations of CBF changes in response to increased CMRO_2_ and CO_2_, and with a study of high altitude acclimatization in which CBF and CMRO_2_ were measured and found to be unchanged from sea level.

### Comparison of the current modeling with earlier modeling of maximum whole body O_2_ metabolism

In a classic study, Wagner (60) modeled the different steps of O_2_ transport from alveolar gas to mitochondria to assess the limiting factors for maximum whole body O_2_ metabolic rate. That work included an equation for O_2_ diffusion from capillary to muscle mitochondria that is mathematically similar to our Eq [6]. However, there is an important difference in the physical interpretation of the parameters: tissue PO_2_ is taken to be mitochondrial PO_2_, and assumed to be near zero for the maximum O_2_ metabolic rate; and the time constant τ is interpreted in terms of the capillary transit time of blood as the ultimate limitation on the diffusion of O_2_ out of the capillary. For our current modeling of O_2_ metabolism in the brain we are not looking at what determines the maximum O_2_ metabolic rate of the brain, but rather how the normal limited range of CMRO_2_ is supported under different conditions. In the context of the tissue O_2_/CO_2_ hypothesis, we specifically are assuming that the mean tissue PO_2_ cannot be too low. A non-zero tissue PO_2_ will drive a back flux of unmetabolized O_2_ from tissue back to the capillary, so that the net O_2_ extraction we focus on here is always less than the unidirectional O_2_ extraction fraction which can be limited by capillary transit time. Put another way, the earlier work essentially assumes that all O_2_ molecules able to leave the capillary are metabolized, and there is no consideration of how O_2_ diffuses in the tissue. Here that tissue diffusion is critical, leading to our interpretation of τ as depending on capillary density.

### Are we focusing on the right tissue PO_2_?

In this work we focused on the average tissue PO_2_. However, the parts of tissue that are most at risk are those with the lowest PO_2_ in the baseline state. In line with that idea, an experimental study from Devor and colleagues (61), in which PO_2_ was measured with high spatial resolution, found that increased CBF maintained the PO_2_ at the locations where PO_2_ was lowest. Our choice to focus on average PO_2_ here was a practical one. Tissue PO_2_ is heterogenous, and we assumed that for most of the empirical data where a PO_2_ is given the value likely corresponds most closely to the average PO_2_. In addition, modeling the lowest value of PO_2_ requires detailed assumptions of the capillary bed geometry and the diffusion characteristics of O_2_ in tissue. For this reason, focusing on the mean value of tissue PO_2_ enables comparison with a wider range of data. For example, in our earlier paper (1) we re-analyzed a previous study of hypoxia (12) that also measured the phosphorylation potential. Using the model we estimated average tissue PO_2_ and found that the phosphorylation potential began to degrade when PO_2_ fell below 12 Torr, in good agreement with the experimental work reviewed by Wilson and colleagues (11). Nevertheless, in light of the paper by Devor and colleagues, the question of where in the tissue O_2_ becomes limiting is an important one that requires further research.

### Summary and conclusions

This work was motivated by considering the entropy available from oxidative metabolism to synthesize ATP. The key idea is that a reduction of the tissue ratio of O_2_ and CO_2_ concentrations could lead to a reduction of the phosphorylation potential, with a cascading effect on all aspects of energy metabolism. Anchoring this thermodynamic hypothesis to empirical measurements suggests that the normal baseline brain tissue O_2_/CO_2_ is only about a factor of 2 higher than the critical level where the phosphorylation potential begins to be impaired. This leads to a general hypothesis that various physiological responses serve to maintain tissue O_2_/CO_2_ in the brain at the baseline level under different conditions. Using a detailed model to calculate tissue O_2_/CO_2_ we found good agreement with the predictions of the hypothesis and reported experimental results in several scenarios: increased oxygen metabolic rate in response to increased neural activity, hypercapnia and hypoxia. For the hypoxia modeling we considered acclimatization to high altitude, adaptation to high altitude in native populations, and the implications of the hypothesis for stroke. The tissue O_2_/CO_2_ hypothesis provides a useful perspective for understanding why many physiological responses under different conditions are useful for preserving brain function, although the mechanisms underlying these functions are not well understood.

## Acknowledgements

The author would like to thank G. Kim Prisk and Susan R. Hopkins for many helpful discussions and suggestions for the manuscript, and also Divya Bolar and David Dubowitz for general discussions of how these ideas might be related to stroke and high altitude physiology.

### Appendix

#### A1. The Krogh cylinder model

The Krogh cylinder model is a classical approach for modeling the exchange of O_2_ with tissue (62). The capillary is treated as a long straight cylinder of radius *a* and length *L*, with O_2_ diffusing out into a larger concentric cylinder with radius *R*. Spatial coordinates are specified in terms of distance along the cylinder *z* (ranging from 0 to *L*) and a radial distance *r* (ranging from *a* to *R* for the extravascular space). The O_2_ concentration in capillary plasma is expressed as an oxygen tension PO_2_ (in Torr), with a solubility α_*O2*_ that converts O_2_ tension to O_2_ concentration (mM).

The standard analysis focuses on a thin disk at *z* in which O_2_ is diffusing from a partial pressure *P*_*C*_(*z*) in the capillary. The rate of O_2_ consumption in the thin disk at *z* is *r*_*O2*_*(z)*, and is assumed to be uniform within the disk. The average of *r*_*O2*_*(z)* over *z* is the mean O_2_ metabolic rate *R*_*O2*_ used in the main text. Radial diffusion of O_2_ is assumed to be described by a diffusion coefficient *D* (units cm^2^/s), and diffusion down the length of the cylinder is neglected. Solution of the diffusion equation for this scenario then provides an expression for tissue oxygen tension as a function of *r* at different positions *z* along the cylinder (62). For brevity we express the tissue PO_2_ as P_T_, and the capillary PO_2_ as P_C_. The distribution is:

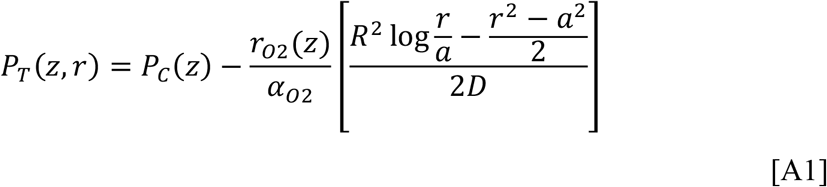

Note that this analysis does not specify how the capillary O_2_ partial pressure PO_2_ varies along the capillary, but only how tissue PO_2_ varies with radial distance for a given capillary PO_2_ profile *P*_*C*_(*z*). To get to the average parameters used in the main text we then average Eq [A1] over *z* and *r* to give:

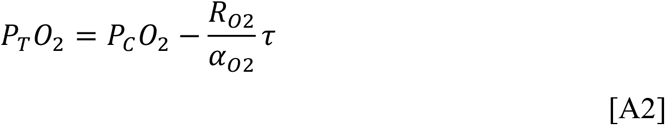

where τ is a time constant defined as the average over *r* of the term in brackets in Eq [A1]. Rearrangement of the terms in Eq [A2] gives Eq [3] in the main text, the primary equation for treating the diffusion of O_2_ in the tissue.

This development in terms of the Krogh model illustrates how Eq [3] arises in a physically plausible way. However, the Krogh cylinder model is likely a poor approximation for the more complex capillary geometry in the brain. For this reason, the expression for τ based on Eq [A1] is not likely to be accurate for the brain. In short, our basic assumption is that the form of Eq [6] is still valid for more complex capillary geometries, but the value of τ needs to be estimated empirically (as described in the main text).

Nevertheless, it is useful to use the Krogh cylinder model to illustrate the physical meaning of τ. The typical time required for an O_2_ molecule to diffuse to the edge of the tissue cylinder is τ_D_=R^2^/2D. The expression in brackets in Eq [A1] also has the form of a distance squared divided by 2*D*, so that τ also has the form of a diffusion time. That is, if we average the expression in the numerator over *r* and call that value *d*^2^, with *d* a distance, τ is then the typical time required for an O_2_ molecule to diffuse that distance *d*. With this interpretation, we can ask how we might expect τ to scale with the capillary volume fraction. For the cylindrical geometry with fixed capillary diameter, as the outer radius *R* increases the tissue volume fraction occupied by the capillary varies as 1/*R*^*2*^, or 1/τ. Based on this reasoning we would expect increased capillary density to lead to a shorter τ.

For CO_2_ clearance, the same diffusion considerations apply, and the equation for average tissue PCO_2_ is:

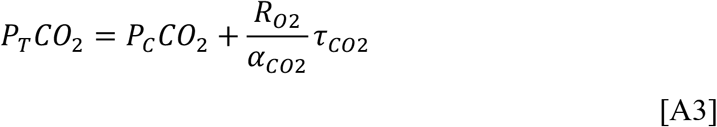

The appropriate τ for CO_2_ is:

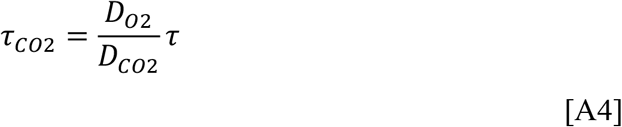

The time constant is only changed by the ratio of the diffusion coefficients. Here we neglect this effect, because the overwhelming effect is the much larger solubility of CO_2_ in Eq [A3] compared with the solubility for O_2_ in Eq [A2]. The result for CO_2_ is that average tissue PCO_2_ is only slightly higher than mean capillary PCO_2_.

#### A2. Transport of O_2_ in blood

In blood, the O_2_ concentration in the form of dissolved gas is a small fraction of the total O_2_ concentration because the majority of O_2_ is bound to hemoglobin. The fractional O_2_ saturation of hemoglobin *S*_O2_ ranges from 0 to 1. As plasma O_2_ diffuses out of the capillary, O_2_ from the hemoglobin is assumed to quickly reach a new equilibrium with the plasma, so that the total O_2_ concentration is:

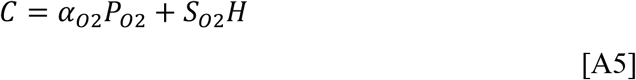

The hemoglobin saturation is assumed to be related to the blood plasma pO_2_ by the Hill equation:

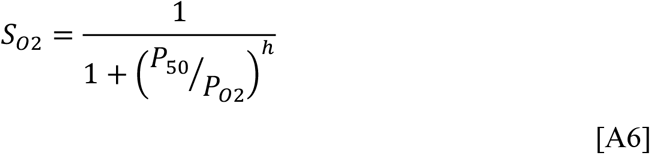

where P_50_ is the P_O2_ value for 50% saturation of hemoglobin, and *h* is the Hill exponent, with values *h*=2.8 and p_50_=27 mmHg often taken as typical values for human blood (63). More recently, Dash and Basingthwaite (59) have shown that the Hill equation, although a simple approximation, can accurately model a wide range of experimental data with the modification that for low P_O2_ values the exponent *h* decreases (i.e., becomes a function of PO_2_). This effect, described as variable cooperativity, increases the O_2_-saturation for the lower PO_2_ values (below ∼30 mmHg). For the variation of *h* at low PO_2_, Dash and Basingthwaite propose:

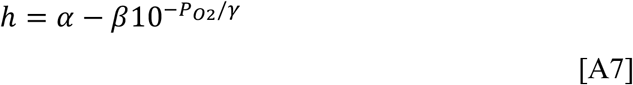

with α=2.82, β=1.2, and γ=29.25 mmHg. This relationship was used in the modeling in the main text.

#### A3. Factors affecting P_50_ of O_2_-hemoglobin binding

Oxygen has a low solubility in water, so efficient transport of O_2_ to tissue requires the majority of O_2_ to be carried bound to hemoglobin. The binding of O_2_ to hemoglobin exhibits a cooperativity effect, in that as more O_2_ is bound, the conformation of the molecule changes in a way that increases the affinity for binding additional O_2_. The value of P_50_, the PO_2_ for 50% saturation of hemoglobin, depends on several factors, including pH, PCO_2_, temperature and the concentration of 2,3-diphosphoglycerate (2,3-DPG, also called 2,3-BPG), a metabolite produced in the red blood cells (59). Higher concentrations of 2,3-DPG reduce the O_2_-binding affinity, producing a rightward shift of the O_2_-hemoglobin binding curve. With prolonged exposure to high altitude the level of 2,3-DPG in blood increases (64).

For the calculations here, we need to consider two aspects of P_50_: 1) the value of P_50_ in the arterial state; and 2) the change in P_50_ as blood moves down the tissue capillary, losing O_2_ and gaining CO_2_ (Bohr effect). In principle, one could model these effects if sufficient information is given about the arterial state of the blood (e.g., see (59) for a recent model). However, for the goal of estimating tissue O_2_/CO_2_ for several populations, sufficient information may not be available, specifically the 2,3-DPG concentration is not routinely reported. Here we take an empirical approach to estimating the P_50_ given other reported parameters of the arterial state, specifically PO_2_ and S_O2_. Given these parameters we estimate the value of P_50_ that would be consistent with the O_2_-hemoglobin modeling in Eqs [A6-A7].

The value of P_50_ varies with species, increasing for smaller animals. In the applications of the modeling we compare the changes needed to preserve the same level of tissue O_2_/CO_2_ for the rat and the human brain. For these estimates we assume P_50_ = 38 Torr for the rat.

For modeling the change in P_50_ from the estimated arterial P_50_ as blood loses O_2_ and gains CO_2_ in the tissue capillary, we assume P_50_ can be modeled as a function of pH:

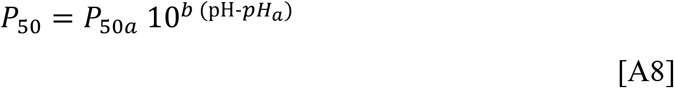

where P_50a_ is the value on the arterial end of the capillary and pH_a_ is the arterial pH (65). The parameter *b* is a negative number called the ‘Bohr factor’. In a recent paper modeling the Bohr and Haldane effects, Malte and Lykkeboe (66) showed that *b* is directly proportional to the number of Bohr groups on each tetramer of hemoglobin. A typical value in human blood is −0.48 (65), which is assumed in the model calculations.

The effects of P_50_ are illustrated in **Figure A1**, showing the rightward shift of the O_2_-hemoglobin saturation curve as P_50_ is increased (**Figure A1.A**, the dashed lines are the high P_50_ case), and the effect of this shift on the estimated average capillary PO_2_ and PCO_2_ (**Figure A1.B**). With the larger value of P_50_ the mean capillary PO_2_ is raised, with no effect on mean capillary PCO_2_.

**Figure A1.**
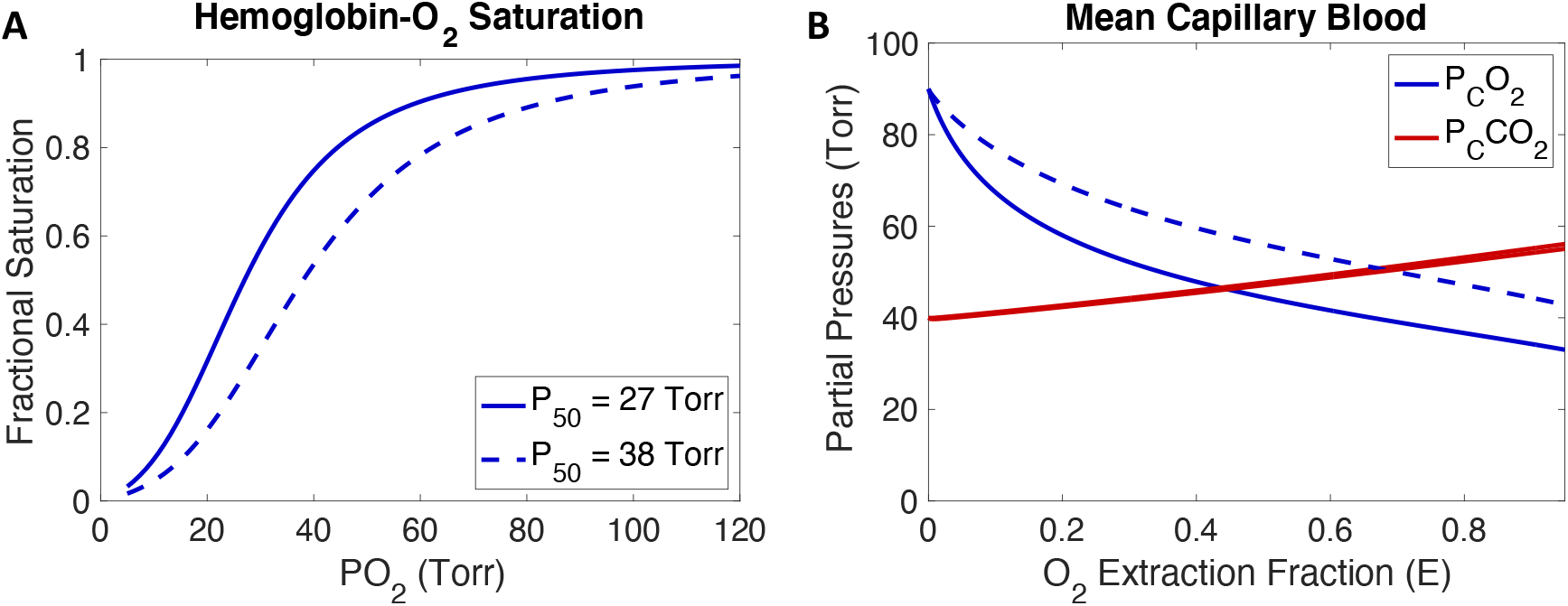
Models used for O_2_ and CO_2_ transport in blood. **A)** O_2_-hemoglobin saturation as a function of PO_2_ for two values of P_50_. **B)** Effect of P_50_ on the calculated average capillary PO_2_ and PCO_2_.

#### A4. Transport of CO_2_ in blood

A model for the transport of CO_2_ is needed in part because the goal is to model changes in tissue PCO_2_ and because additional CO_2_ carried in blood will affect the pH that in turn affects P_50_ and the average capillary PO_2_. Carbon dioxide diffuses from tissue into blood as dissolved gas, but once in the blood the majority is converted to bicarbonate ions (HCO_3_^-^) in plasma, catalyzed by the enzyme carbonic anhydrase. In addition, CO_2_ binds to hemoglobin to form carbamino compounds, which are a significant carrier of CO_2_ in venous blood. This complexity of CO_2_ carriage in blood makes the relationship between total CO_2_ (T_CO2_) and PCO_2_ somewhat complicated, depending on hemoglobin content (H), pH, and hemoglobin O_2_-saturation S_O2_.

We assume the total CO_2_ concentration in blood (in mM) is (59):

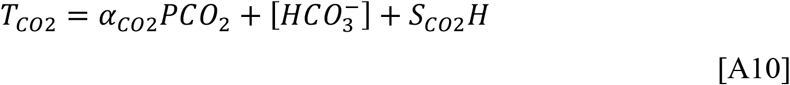

The fractional CO_2_ saturation of hemoglobin is S_CO2_, and *H* is the hemoglobin concentration as defined in Section 3.1. We assume that the bicarbonate concentration [HCO_3_^-^] is always in equilibrium with the PCO_2_, described by the Henderson-Hasselbalch equation:

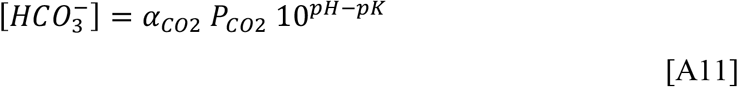

with pK=6.091.

The remaining modeling question for CO_2_ transport is how to approximate the CO_2_ saturation of hemoglobin, which depends on PCO_2_, PO_2_ and pH. Here we used a simplified expression created to approximately match the detailed modeling of Dash and Basingthwaite (59) (based on their Figure 4). (In that work S_CO2_ is calculated as a function of changes in pH_RBC_, the red blood cell pH, and for our calculations we converted to plasma pH changes with the factor of 0.77 (67)). The approximation is then:

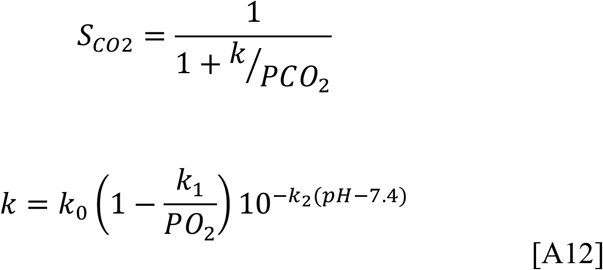

For partial pressures in Torr, the constants are: k_0_=900; k_1_=14; and k_2_=1.16. Note that the form of Eq [A12] is simply a convenient mathematical form, rather than one based on the underlying physics, that approximates the curves in Dash and Basingthwaite for PCO_2_ up to 100 mmHg and pH up to 8.0.

#### A5. pH changes in the tissue capillary as O_2_ and CO_2_ are exchanged

To calculate the changes in capillary PO_2_ and PCO_2_ as blood moves down the capillary we need to consider how the various blood parameters change in an interconnected way as O_2_ is exchanged one for one with CO_2_ (i.e., a respiratory quotient of 1.0 consistent with oxidative metabolism of glucose in the brain). Blood pH changes down the length of the capillary as CO_2_ is added, equivalent to adding an acid, but the buffering in the blood also changes with S_O2_ through the effects of hemoglobin (deoxy-hemoglobin is a more effective buffer than oxy-hemoglobin). The pH buffering of blood is described as a buffer line in a Davenport plot of bicarbonate vs pH. Our assumptions here are based on the recent modeling work of O’Neill and Robbins (68). Using their model, they found slopes of the buffer lines of oxy-hemoglobin and deoxy-hemoglobin of −25.6 and −24.7 mM/pH unit, respectively, with a separation between the buffer lines of 0.084 pH units for a bicarbonate concentration of 25 mM. They found good agreement with experimental values of −28.0, −27.8 and 0.084 for these three values. To simplify the equations, we assumed equal slopes for the two curves. For the magnitude of the slope we assumed the average of the empirical and theoretical values, −27 mM/pH. The pH separation of the two curves was assumed to be 0.084 for [HCO_3_^-^]=25 mM.

With these assumptions, and assuming a linear shift between the oxy-hemoglobin and deoxy-hemoglobin buffer lines as S_O2_ is reduced from full oxygenation, the buffer curve relating [HCO_3_^-^] to pH is modeled as:

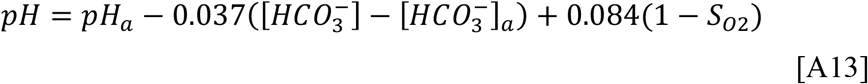

#### A6. Application of the model

The key application of the modeling described in **Sections A2-A5** is to calculate the average capillary PO_2_ and PCO_2_ for a given arterial state and given values of blood flow and O_2_ metabolism. To do this, we step through values of the extraction fraction *E*, which determines the amount of O_2_ removed with an equal amount of CO_2_ added to the blood. We then calculate what the venous values of PO_2_ and PCO_2_ would be for that value of *E*. The average values of capillary PO_2_ and PCO_2_ for a given net extraction *E* are then estimated as the average of the venous values for all extraction fractions less than *E*.

For a given value of *E*, removing the corresponding amount of O_2_ and adding an equal amount of CO_2_ changes both total O_2_ and total CO_2_ by an increment ΔTO_2_. For initial values TO_2_(0) and TCO_2_(0), the new total O_2_ and CO_2_ are then:

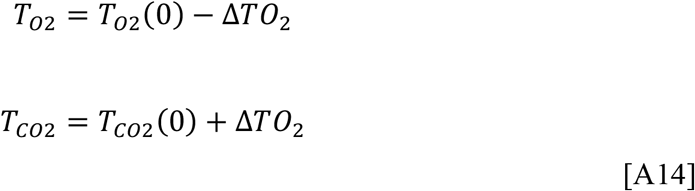

(Note that the one-for-one exchange of O_2_ for CO_2_ reflects the assumption that the respiratory quotient has a value of one for the brain.) The new TCO_2_ will distribute between dissolved gas, bicarbonate ions, and carbamino-bound CO_2_, creating new values of PCO_2_, [HCO_3_^-^] and pH. The goal is to find a consistent solution to the equilibrium equations for CO_2_, Eqs [A10-A12].

To find this solution, we do an iterative calculation for each addition of ΔTO_2_. The bicarbonate value [HCO_3_^-^] is incremented by a small amount, and a new pH calculated based on the buffering curve (Eq [A13]). Then a new PCO_2_ value is calculated from the bicarbonate/PCO_2_ equilibrium relation (Eq [A11]). The Bohr effect is then modified by updating P_50_ for O_2_ binding to hemoglobin using the new pH value (Eqs [A8-A9]), and a new PO_2_ is calculated for the new total O_2_ (Eqs [A5-A7], requiring a numerical inversion). Using the new value of PO_2_, with the new values of pH and PCO_2_, the carbamino-bound CO_2_ is calculated with Eq [A12]. Finally, the new total CO_2_ is calculated using all the new values. This process is repeated, updating the bicarbonate concentration with each step, until the total CO_2_ calculated with Eq [A10] reaches the desired level for the increase by ΔTO_2_ defined by Eq [A14].

